# The extracellular morphology and behaviour of the last common ancestor of animals and choanoflagellates

**DOI:** 10.64898/2026.11.12.765648

**Authors:** Margarita Skamnelou, Cédric Berney, Guifré Torruella, Vasily Zlatogursky, Miguel Ángel Fernández Alcón, Luis Javier Galindo, Yana Eglit, Daryna Zavadska, Giacomo Mutti, Juliette Mathieu, Patrick J. Keeling, David Moreira, Purificación López-García, Daniel J. Richter

**Affiliations:** Institut de Biologia Evolutiva (CSIC-Universitat Pompeu Fabra), Barcelona, Spain; Department of Organismal Biology, Uppsala University, Uppsala, Sweden; Department of Botany, University of British Columbia, Faculty of Science,Vancouver, BC, V6T 1Z4, Canada; Institute of Water Research, University of Granada, Granada, Spain; Department of Ecology, University of Granada, Campus Fuentenueva, Granada, Spain; Department of Biology, and Centre for Comparative Genomics and Evolutionary Bioinformatics, Dalhousie University, Halifax, NS, Canada; Institut de Recerca Biomèdica, Barcelona, Catalonia, Spain; Barcelona Supercomputing Centre (BSC-CNS), Barcelona 08034, Spain; Centre for Interdisciplinary Research in Biology, Collège de France. CNRS UMR7241, Paris, France; Unité d’Ecologie Société et Evolution, CNRS, Université Paris-Saclay, AgroParisTech, Gif sur Yvette 91190, France

**Keywords:** choanoflagellate, theca, lorica, glycocalyx, colony, filopodia, animal evolution, transcriptome, phylogenetics, taxonomy

## Abstract

Choanoflagellates, the closest known living relatives of animals, provide a powerful comparative system for investigating the evolution of extracellular morphology and cell behaviour on the animal stem lineage. Although all choanoflagellates share a conserved cellular architecture including a flagellum and collar of microvilli, they display a wide diversity of extracellular structures and behaviours throughout the life cycle. Their extracellular features, historically used for species identification, have long been suspected to reflect a mixture of evolutionary inheritance and ecological plasticity. Yet, the majority of choanoflagellate diversity remains undescribed, and the extent to which their extracellular traits map onto their phylogeny has remained unclear, impeding comparative reconstructions of the evolutionary history of extracellular structures with the earliest animals. Here, we integrate super-resolution fluorescence imaging, electron microscopy and transcriptomic analysis to characterize the extracellular structures and behaviours across both currently described lineages of choanoflagellates (Craspedida and Acanthoecida), by studying pairs of closely related species in conjunction with novel isolates representing previously unexplored clades. Our results reveal strong conservation of extracellular morphology between close relatives, whereas deeper evolutionary distances uncover unexpected innovations, such as sheath-like layers, bipartite thecae, crawling stages and atypical inverted division patterns. Furthermore, the inclusion of early-branching taxa sister to the loricates supports the hypothesis that the silicified extracellular lorica is a derived trait rather than an ancestral feature. Together, our findings suggest that the last choanoflagellate common ancestor possessed an organic non-silicified extracellular covering composed of one or more layers and that it was capable of both swimming and crawling behaviours, refining the cellular context from which animals evolved.

## Introduction

Choanoflagellates are unicellular or colonial heterotrophic protists that inhabit a diverse range of ecological habitats from pole to pole, including marine, brackish, freshwater, soil, and even certain extreme environments (Richter and Nitsche, 2017; Schiwitza et al., 2018). Their conserved morphology consists of a cell body with an apical motile flagellum surrounded by a collar of microvilli (Hibberd, 1975; Karpov, 2016; Karpov and Leadbeater, 1998; Leadbeater, 2014; Richter and Nitsche, 2017). The movement of the single flagellum generates fluid flow, facilitating the delivery of bacterial prey into the feeding collar for subsequent phagocytosis (Dayel and King, 2014; Leadbeater and Morton, 1974; Pettitt et al., 2002). Within marine and freshwater ecosystems, this particular trait makes them competent filter feeders, thereby exerting a strong ecological impact within microbial food webs (Leadbeater, 2014).

Beyond their ecological importance, choanoflagellates hold a unique phylogenetic position. In addition to their apparent structural similarity to the choanocytes in sponges (Kent, 1882), phylogenetic studies place choanoflagellates as the closest living relatives of metazoans (King et al., 2008; King and Carroll, 2001; Ruiz-Trillo et al., 2008; Snell et al., 2001). This makes them uniquely informative both for reconstructing features of the last common ancestor of choanoflagellates and animals and for distinguishing ancestral traits from metazoan-specific innovations. For example, genomic and transcriptomic studies have revealed the existence of gene families in choanoflagellates that were previously thought to have been animal-specific (Fairclough et al., 2013; Gao et al., 2024; Nichols et al., 2012; Richter et al., 2018; Ros-Rocher et al., 2021; Ruiz-Trillo et al., 2023), enabling the study of the pre-metazoan evolution of these gene families.

Thus far, a small number of choanoflagellate species have emerged as valuable model organisms, but studies utilizing environmental sequences have revealed that the diversity within choanoflagellates remains largely uncharacterized (Del Campo and Massana, 2011; Del Campo and Ruiz-Trillo, 2013; Venter et al., 2018; Weber et al., 2012). From a taxonomy viewpoint, choanoflagellates are divided into two orders: Craspedida and Acanthoecida (Carr et al., 2017, 2008). However, phylogenetic analysis of environmental 18S rRNA gene sequences indicates a vast diversity of uncharacterized species both within and outside these two orders (Del Campo and Ruiz-Trillo, 2013; Jiménez-Valadez et al., 2026; López-Escardó et al., 2019). In contrast to the consistent morphology of the choanoflagellate cell itself (with a single flagellum surrounded by a collar of microvilli), there is considerable variation in choanoflagellate extracellular structures (Carr et al., 2008; Nitsche et al., 2011). Acanthoecid species construct an external silicified covering called a lorica, whereas craspedids may produce an organic rigid extracellular theca or a flexible sheath which, in some species, has been shown to contain carbohydrates (Carr et al., 2017; Dayel et al., 2011; Garcia De Las Bayonas et al., 2026; Leadbeater and Morton, 1974). Choanoflagellates have also been found to secrete additional extracellular material during colony formation (including the extracellular matrix protein Rosetteless), and during other life history stages such as the cystic stage, in which the composition of the extracellular material remains unknown (Leadbeater and Karpov, 2000; Levin et al., 2014; Stoupin et al., 2012). Beyond these morphological differences, the biochemical composition of most extracellular structures remains poorly described, as do their functions, which may include surface attachment, connection of cells within colonies, mediation of gamete adhesion and fusion during sexual reproduction, protection, flotation, hydrodynamic efficiency, or a possible association with iron uptake, as recently demonstrated in thecate cells (Dayel et al., 2011; Garcia De Las Bayonas et al., 2026; Leadbeater, 2014; Leon et al., 2025; Levin et al., 2014). The above cases suggest that, if these extracellular features were present in the last common ancestor of animals and choanoflagellates (i.e., Choanozoa), the evolutionary roots of animal extracellular organization may have predated animal origin. Recent methodological advances, such as the application of biochemical staining and super-resolution microscopy, now enable more refined characterization of extracellular structures, opening the way to a deeper understanding of their diversity, composition and evolutionary history (Colin et al., 2017; Huff, 2015). Such advances may help determine which extracellular features represent ancestral characteristics of the last common ancestor of animals and choanoflagellates, which are lineage-specific innovations, and clarify the potential evolutionary relationships between the extracellular material of choanoflagellates and the extracellular systems associated with animal multicellularity.

To address these questions, we used the fluorescence method e-HCFM (environmental High Content Fluorescence Microscopy) (Colin et al., 2017) to visualize extracellular structures, performed 3D cell reconstructions and compared the morphologies of pairs of closely related species or groups, including both newly isolated and previously described strains sampled from diverse clades of the choanoflagellate tree. By coupling fluorescence and super-resolution imaging with electron microscopy and time-lapse observations, we identify strong conservation of extracellular morphologies among closely related species, contrasted with marked divergence across distant lineages. Our analyses also reveal unexpected features, including sheath-like coverings that closely envelop the cell body, two-parted tubular thecae capable of partial shedding, and crawling stages that emerge from the theca after cell division. Notably, we report for the first time an inverted mode of division within thecate species. In addition, by characterizing novel isolates from the previously uncultured FRESCHO-3 clade (Del Campo and Ruiz-Trillo, 2013), we provide evidence that the lorica is a derived innovation within Acanthoecida rather than an ancestral feature of choanoflagellates. Taken together, these results expand the known diversity of choanoflagellates, provide new insights into their biology, reveal evolutionary patterns linking extracellular morphology with phylogeny and enable us to propose informed hypotheses about the extracellular morphology of both the choanoflagellate last common ancestor, and the ancestor they share with animals.

## Results

### Phylogenetic framework for comparative morphological analysis

To expand available phylogenetic diversity for our comparisons, we complemented previously described strains (*Salpingoeca rosetta*, *Stagondoeca pyriformis* and *Salpingoeca tuba*) with five novel isolates, including representatives of both known and previously underexplored clades: *Hartaetosiga kampana* sp. nov. strain BEAP0001, *Oinochoeca dipylis* gen. et sp. nov. strain BEAP0002, *Salpingoeca kylix* sp. nov. strain BEAP0192, *Daedalina diplosolina* gen. et sp. nov. strain BEAP0272 and *Hippolytella eleftheri* gen. et sp. nov. strain BEAP0279. We sequenced and assembled high quality and nearly complete transcriptomes, with BUSCO (Manni et al., 2021; Simão et al., 2015) completeness scores of 90.2% for *Hartaetosiga kampana*, 94.2% for *O. dipylis,* 92.6% for *Salpingoeca kylix*, 93.3% for *D. diplosolina*, and 87.5% for *Hippolytella eleftheri*.

Phylogenomic analysis was first conducted using PhyloFisher (Tice et al., 2021) to identify orthologous markers, followed by maximum likelihood inference with IQ-TREE (Wong et al., 2026). This analysis included the five newly assembled transcriptomes along with 26 additional choanoflagellate species and 22 outgroup holozoans (9 protists and 13 metazoans) (**Figure 1A**). To increase taxon coverage, particularly for species lacking transcriptomes, such as *Stagondoeca pyriformis, Salpingoeca tuba*, and FN-2015, a species representing the FRESCHO-3 group, a phylogenetic tree was constructed using concatenated 18S, 5.8S, and 28S rRNA gene sequences from 52 choanoflagellate taxa and 10 outgroup holozoans (2 protists and 8 metazoans) (**Supplementary Figure 1**).

**Figure 1:**
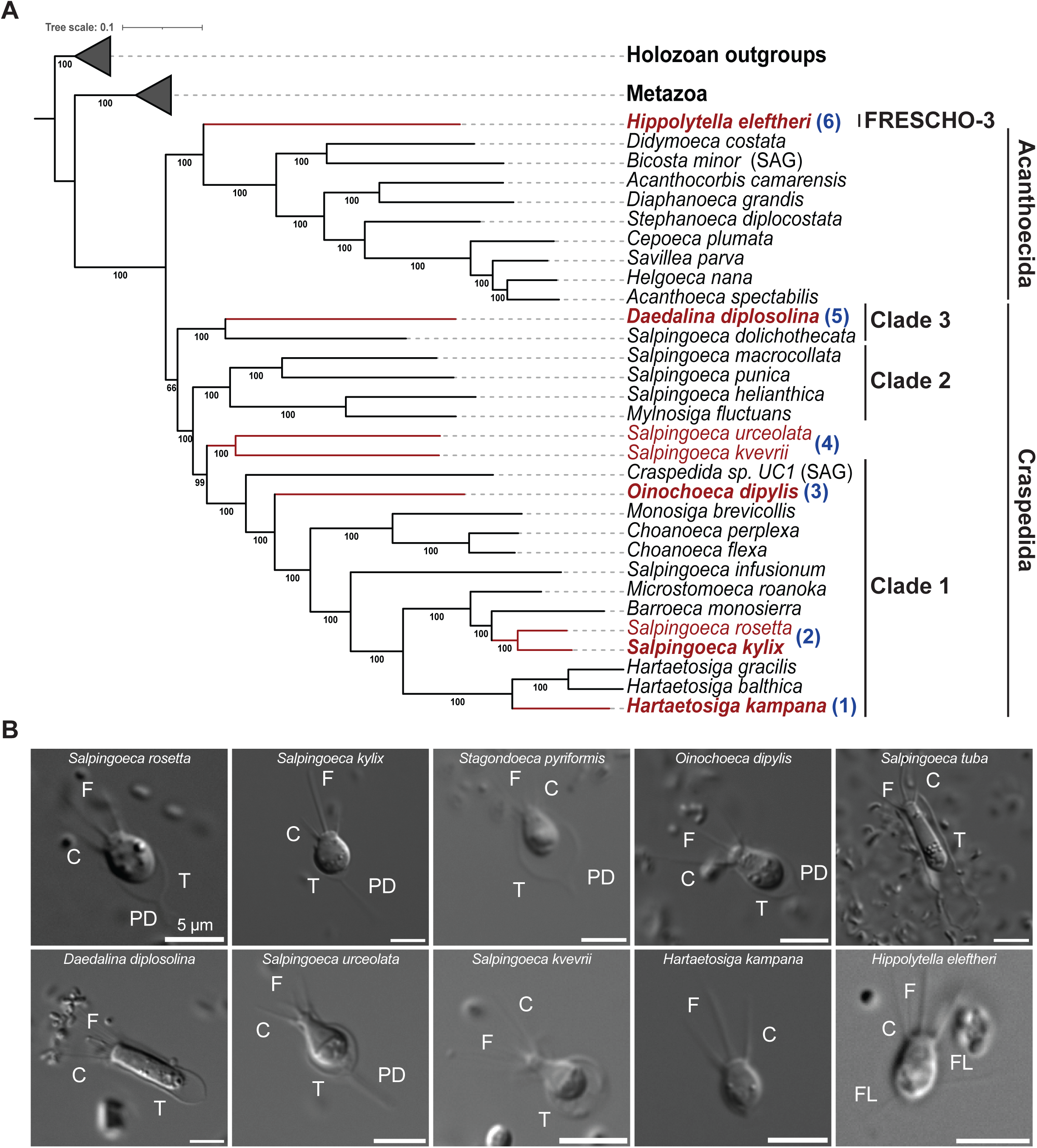
Phylogenetic reconstruction and initial morphological assessment of selected choanoflagellate species. **A)** Phylogenomic reconstruction, based on transcriptome data from five newly isolated species shown in bold type (*Hartaetosiga kampana*, *Salpingoeca kylix*, *Daedalina diplosolina*, *Oinochoeca dipylis* and *Hippolytella eleftheri*) and 26 additional choanoflagellates, alongside 21 outgroup holozoans (13 Metazoa, 9 protists). Outgroups collapsed for clarity. The species whose extracellular structures were visualized and compared in this study are highlighted in red. Orthologous markers were identified using PhyloFisher (Tice et al. 2021), followed by maximum likelihood inference using IQ-TREE (Wong et al., 2026). Bootstrap values are shown below internal branches. *H. kampana* is within the genus *Hartaetosiga*, *S. kylix* branches with *Salpingoeca rosetta*, *Salpingoeca kvevrii* and *Salpingoeca urceolata* form a strongly supported clade, *O. dipylis* is found in a distinct, well-supported lineage in Clade 1, *D. diplosolina* branches with *Salpingoeca dolichothecata* in Clade 3, and *H. eleftheri* is positioned as a sister to loricates (Acanthoecida). “SAG” label indicates markers obtained from single-cell amplified genomes. The blue labels indicate the order of morphological comparisons described in the text. A phylogenetic tree with increased taxon sampling, reconstructed based on the ribosomal RNA gene locus, is shown in Supplementary Figure 1. **B)** Differential interference contrast (DIC) microscopy images showing representative morphology of the analysed species in the presence of bacterial prey. All species displayed the conserved ovoid cell body typical of the Choanoflagellatea class, characterized by a single flagellum (F) surrounded by a collar of microvilli (C). Most species (except *Hartaetosiga kampana* and *Hippolytella eleftheri*) can exhibit an extracellular theca (T), in many cases supported by an extended pedicel (PD). Filopodia (FL) were also observed in some taxa. Scale bar: 5 μm.

*Hartaetosiga kampana* was recovered within the genus *Hartaetosiga* with strong support, while *Salpingoeca kylix* was closely related to *Salpingoeca rosetta*. *Oinochoeca dipylis* formed a distinct, well-supported lineage representing a novel genus, sister to *Stagondoeca pyriformis*. *Daedalina diplosolina* was placed within Clade 3 as a sister genus to the clade containing *Salpingoeca dolichothecata* and *Salpingoeca tuba*. The previously suggested close relationship between *Salpingoeca kvevrii* and *Salpingoeca urceolata* was also confirmed. Notably, *Hippolytella eleftheri* and the choanoflagellate FN-2015 together formed a strongly-supported sister group to the Acanthoecida.

Based on the phylogenetic relationships among our newly isolated and previously sequenced choanoflagellates, we defined six groups for comparison, each consisting of closely related taxa sampled from across choanoflagellate diversity: (1) *Hartaetosiga kampana*, a novel *Hartaetosiga* species, versus other members of the *Hartaetosiga* genus, (2) *Salpingoeca kylix* versus *Salpingoeca rosetta*, (3) *Oinochoeca dipylis*, representing a novel genus, versus *Stagondoeca pyriformis*, (4) the flask-shaped species *Salpingoeca kvevrii* and *S. urceolata*, (5) *Daedalina diplosolina*, a newly isolated Clade 3 genus and species, versus other tubular thecate choanoflagellates from Clade 3, and (6) *Hippolytella eleftheri*, one of the first cultured representatives of the FRESCHO-3 clade, compared with acanthoecids to investigate lorica evolution. All selected species were first examined by differential interference contrast (DIC) microscopy (**Figure 1B**). Consistent with the previously described conserved morphology of the Choanoflagellatea class (Karpov and Leadbeater, 1998; Richter and Nitsche, 2017), cells from all examined species have a characteristic single flagellum surrounded by a collar of microvilli. In all the species except *Hartaetosiga kampana* and *Hippolytella eleftheri*, the presence of an extracellular structure resembling a theca was observed, often distinguished by an extended pedicel.

### 1. The *Hartaetosiga* genus is characterized by highly conserved extracellular morphology

The novel choanoflagellate species *Hartaetosiga kampana* was isolated from the shallow marine lagoon Malo Jezero on Mljet Island, Croatia. *H. kampana* cells were stained using markers for DNA (Hoechst), the collar of microvilli (phalloidin) and the flagellum (β-tubulin) and imaged by using super-resolution microscopy. The cells are characterized by an elongated body shape and a collar of microvilli that widens as it extends away from the cell (**Figure 1B**, **Figure 2A**).

**Figure 2:**
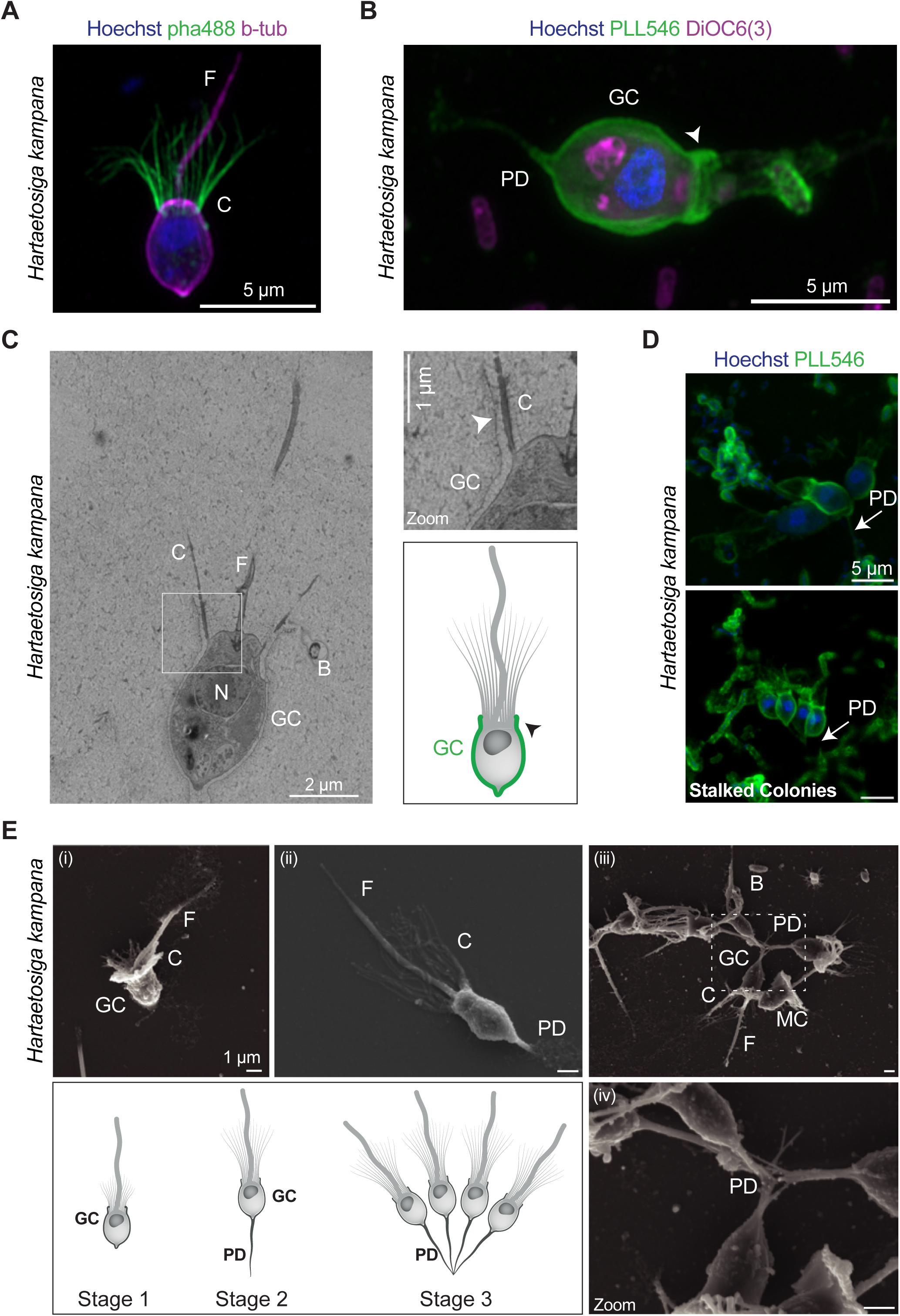
Morphology, extracellular properties and life cycle of *Hartaetosiga kampana*, in the presence of co-cultured bacterial prey. C: collar, F: flagellum, GC: glycocalyx, N: nucleus, PD: pedicel. Arrowhead indicates the glycocalyx coat covering part of the collar of microvilli. **A-B)** Super-resolution confocal microscopy of individual *H. kampana* cells stained with the indicated markers. Hoechst (blue): DNA content, pha488 (green): F-actin, PLL546 (green): extracellular structures, DiOC6(3) (magenta): Cell body membranes, and β-tub (magenta): flagellum. Scale bar: 5 μm. **C)** Transmission Electron Microscopy (TEM) section and drawing of *H. kampana* showing a thin, and close-fitting glycocalyx coat that covers a part of the collar of microvilli (highlighted by arrowhead). Scale bar: 2 μm and 1 μm (in zoom). **D)** Confocal microscopy of stalked colonies of *H. kampana* isolate stained as in B (Hoechst, PLL546). Scale bar: 5μm. **E)** Scanning electron microscopy (SEM) images and drawings of *H. kampana* at distinct stages. (i) Motile cell with short collar, and flagellum. (ii) Attached cell via pedicel with long collar, and flagellum. (iii) Colonial cells with long collar, flagellum and pedicel. White box indicates connections among cells in a stalked colony and is shown in higher magnification in (iv). Scale bar: 1μm.

To gain further insight into the extracellular architecture, we applied the e-HCFM fluorescence staining method (Colin et al., 2017), which allows simultaneous visualization of DNA (Hoechst stain), membranes and hydrophobic bodies (DiOC6(3) stain), and extracellular components (α-poly-L-lysine conjugated to Alexa Fluor 546, heretofore PLL546 stain). This staining approach was chosen because poly-L-lysine (PLL) binds to cell surfaces, allowing us to label extracellular structures whose exact composition remains unknown. The cell surface of *H. kampana* is surrounded by a thin extracellular coat that is referred to in the literature as a glycocalyx (Carr et al., 2017, 2008; Leadbeater and Morton, 1974; Schiwitza et al., 2023; Wylezich et al., 2012) (**Figure 2B**). This coat can extend into a pedicel and partially covers the collar of microvilli (**Figure 2B**, white arrowhead). Transmission electron microscopy (TEM) further confirmed that the glycocalyx forms a thin, closely-fitting layer around the cell body and extends over part of the collar (**Figure 2C**). The presence and structural features of the glycocalyx appear to be conserved within the *Hartaetosiga* genus, namely: its close-fitting nature, ability to form a pedicel and partial coverage of the collar (Carr et al., 2017, 2008; Leadbeater and Morton, 1974; Schiwitza et al., 2023; Wylezich et al., 2012).

In addition to these structural features, *H. kampana* displays a life cycle consistent with that of other *Hartaetosiga* species, comprising three different life history stages: (1) motile cells with a single flagellum surrounded by a collar composed of short microvilli (**Figure 2E-i**), (2) attached cells with an extruded pedicel, a flagellum and a collar composed of long microvilli (**Figure 2E-ii**), and (3) multicellular colonies (**Figure 2D** and **Figure 2E-iii-iv**), attached to the culture flask surface through extended pedicels joined at their basal extremities and collars of long microvilli. The microvillar collar of non-attached, potentially motile cells is relatively short in comparison to the longer microvilli composing the collars of attached cells. Previous studies have proposed that such alterations in the length of the flagellum and microvilli can confer advantages in swimming speed and feeding capability (Nguyen et al., 2019).

### 2. Conserved extracellular features between *Salpingoeca rosetta* and *Salpingoeca kylix*

The novel choanoflagellate species *Salpingoeca kylix* was isolated from a salt evaporation pond in Eilat, Israel and is closely related to *Salpingoeca rosetta* (Clade 1) (**Figure 1A and Supplementary Figure 1**). Super-resolution microscopy confirmed the presence of an organic extracellular covering, known as a theca, in both *Salpingoeca* species (**Figure 3A**), consistent with the observations made using DIC microscopy and in line with the known characteristics of *S. rosetta* (Dayel et al., 2011) (**Figure 1B**). We compared thecate cells and empty thecae, stained with PLL546, of both *Salpingoeca* species with super-resolution microscopy, followed by 3D reconstructions using either serial sections (**Figure 3B-i**) or surface analysis (**Figure 3B-ii-iii**). This analysis revealed that both species exhibited a cup-shaped theca, covering the bottom end of the cell. Interestingly, the thecate cells were characterized by an additional stained layer that partially covered the collar of microvilli and the cell body, in a close-fitting manner (**Figure 3B**, indicated by a white asterisk). Confocal fluorescence imaging demonstrated that the additional coat remained associated with the cell even after the theca was shed (**Supplementary Figure 2A**). A similar structure was previously described in *S. rosetta* as a “skirt” (Dayel and King, 2014).

**Figure 3:**
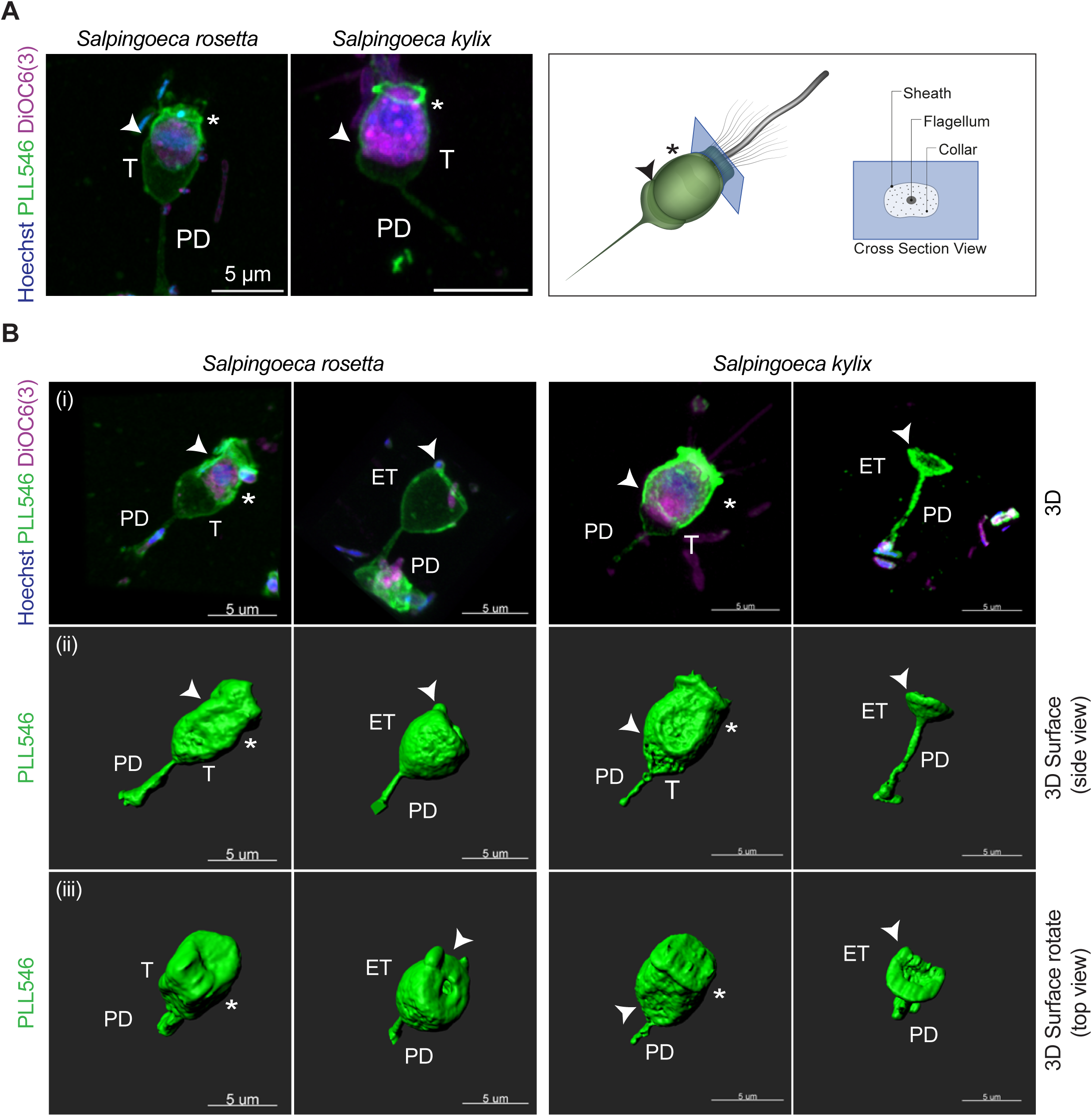
Comparative analysis of theca morphology in *Salpingoeca rosetta* and *Salpingoeca kylix*, together with co-cultured bacterial prey, using super-resolution fluorescence microscopy of cells stained with Hoechst (blue, DNA), PLL546 (green, extracellular structures) and DiOC6(3) (magenta, cell membranes). Both species exhibit a theca (T) and a pedicel (PD). **A)** Super-resolution images of individual *Salpingoeca* cells. White arrowheads indicate the distal end of the theca and asterisks mark an additional sheath-like layer surrounding the cell. Scale bar: 5 μm. **B)** 3D reconstructions from super-resolution confocal images of thecate cells and empty thecae (ET). For each species, three visualizations are shown: (i) Overlay of all three fluorescence channels, (ii) 3D surface reconstruction of PLL546 staining, and (iii) Rotated view of panel (ii). White arrowheads indicate the distal end of the theca and asterisks mark an additional sheath-like layer surrounding the cell. Both *S. rosetta* and *S. kylix* display a cup-shaped theca extending into a pedicel, as well as a flexible outer coat partially covering the collar and cell body.

In addition, scanning electron microscopy (SEM) of *S. kylix* identified three different life forms, similar to those reported for *S. rosetta*: (1) cells with an additional sheath without theca and pedicel (likely slow swimmers, *sensu* Dayel et al., 2011 (Dayel et al., 2011)) (**Supplementary Figure 2B-i-ii**), (2) attached cells with an additional external sheath that are enclosed in a cup-shaped theca with an extended pedicel characterized by a collar of microvilli (**Supplementary Figure 2B-iii-iv**), (3) bean-shaped cells with a short collar, long flagellum and extended filopodia (likely fast swimmers, *sensu* Dayel et al., 2011 (Dayel et al., 2011)) (data not shown)

### 3. Conserved theca shape between Stagondoeca pyriformis and Oinochoeca dipylis

The novel choanoflagellate genus and species *Oinochoeca dipylis* was isolated from Janubio beach in Lanzarote, Spain. A closely related species (**Figure 1A)**, *Stagondoeca pyriformis*, originally described by Carr et al. (2017) (Carr et al., 2017), was re-isolated during this study from a beach in Lido di Pomposa, Ferrara, Italy. Both species exhibited capsule-shaped thecae with three distinct curved lobes (labelled 1, 2, and 3 in **Figure 4** and also seen in **Supplementary Figure 3A**). In *O. dipylis*, however, all empty thecae suitable for 3D surface reconstruction lacked the third lobe (**Figure 4A**, second row), likely due to mechanical damage caused by cells exiting the theca. Occasional absence of the third lobe was also noted in *S. pyriformis*, likely for similar reasons. Nonetheless, intact PLL546-stained empty thecae of *O. dipylis* (**Figure 4C-i**) support the notion that the three-lobed configuration is typical in both species.

**Figure 4:**
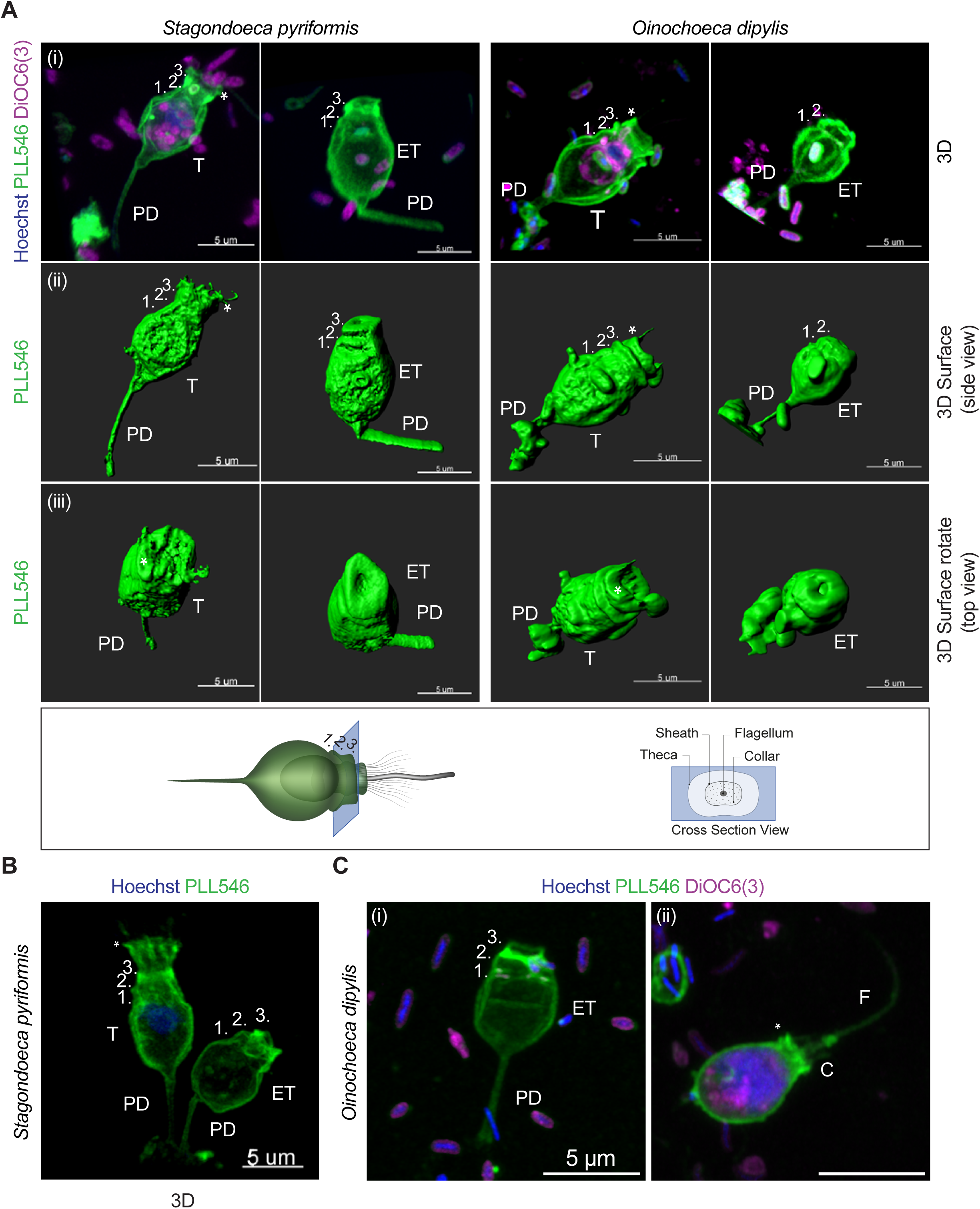
Extracellular structure comparison between *Stagondoeca pyriformis* and *Oinochoeca dipylis*, stained with Hoechst (blue): DNA content, PLL546 (green): extracellular structures, DiOC₆(3) (magenta): cell body membranes. T: theca, ET: empty theca, PD: pedicel. Scale bar: 5 μm. **A)** D super-resolution confocal microscopy reconstructions of thecate cells and empty thecae from each species. Three visualizations are shown: (i) Overlay of all three fluorescence channels, (ii) 3D surface reconstruction of PLL546 staining, and (iii) Rotated view of panel (ii). Both species exhibit a capsule-shaped theca with three curved lobes (indicated by numbers 1, 2, and 3). Asterisk highlights an additional extracellular sheath partially covering the collar of microvilli. In the surface analysis (ii-iii), the PLL546-stained sheath (asterisk) is more distinct in *O. dipylis* than *S. pyriformis*. In *O. dipylis*, empty thecae suitable for 3D surface reconstruction lack the third lobe (second row), likely due to mechanical breakage during cell exit. **B)** A thecate cell and an empty theca of *S. pyriformis* stained with PLL546 and Hoechst. The additional extracellular sheath is clearly visible (asterisk), and the three lobes of the theca are marked by numbers 1, 2 and 3. **C)** An empty theca (i) and a cell lacking a theca (ii) in *O. dipylis*. The empty theca displays all three lobes in their intact configuration, highlighting structural similarity with *S. pyriformis*. The cell without a theca retains an extracellular sheath (asterisk) closely envelopes both the cell body and the collar.

In both species, thecate cells often displayed an additional extracellular sheath, also stained by PLL546, closer to the cell body than the theca and partially covering the collar of microvilli (**Figure 4A-B**, indicated by an asterisk). In *S. pyriformis*, the extracellular sheath was less apparent in surface reconstructions, likely due to its overlapping position with the thecal neck (**Figure 4A-ii-iii**). However, its presence is visible by 3D reconstruction fluorescence imaging in both species (**Figure 4A-i, B**). Notably, in *O. dipylis*, cells lacking a theca but retaining the sheath were also observed (**Figure 4C-ii**). In these cases, the sheath appears to closely envelop not only the collar but the entire cell body.

Further analysis of *O. dipylis* thecate cells by TEM supported the presence of this additional extracellular sheath (**Supplementary Figure 3A**). Moreover, this analysis showed that the theca is thin and separated from the cell body. Interestingly, unlike the observations made during fluorescence imaging (**Figure 4C-ii**), the sheath was not visibly covering the entire cell body in the TEM sections. This difference suggests that the sheath may either be extremely closely associated with the membrane, making it difficult to resolve by TEM, or too delicate to withstand TEM sample preparation and fixation.

Overall, the theca morphology observed in the re-isolated *S. pyriformis* strain BEAP0306 differs from its original characterization as the type species of the *Stagondoeca* genus via DIC microscopy, in which the theca was described as having a droplet-like shape without a neck, enclosing the entire cell body (Carr et al., 2017). Therefore we emend the diagnosis of *Stagondoeca* to update it based on higher-resolution observations.

### 4. Conserved extracellular morphology between *Salpingoeca urceolata* and *Salpingoeca kvevrii*

We next expanded the comparative study to include craspedids outside Clade 1. To this end, we obtained two previously cultured, characterized and closely related species, *Salpingoeca kvevrii* and *Salpingoeca urceolata* (Carr et al., 2017; Richter et al., 2018).

Super-resolution microscopy confirmed the presence of the theca in both species (**Figure 5A**). This finding is consistent with DIC microscopy observations and aligns with previous characterizations of the two species (Carr et al., 2017) (**Figure 1B**). Both species exhibited a characteristic flask-shaped theca with a distinct neck (**Figure 5**, indicated by arrowhead), consistent with earlier descriptions. Notably, the theca of *S. urceolata* can extend into a pedicel, a feature that has not been observed in *S. kvevrii*.

**Figure 5:**
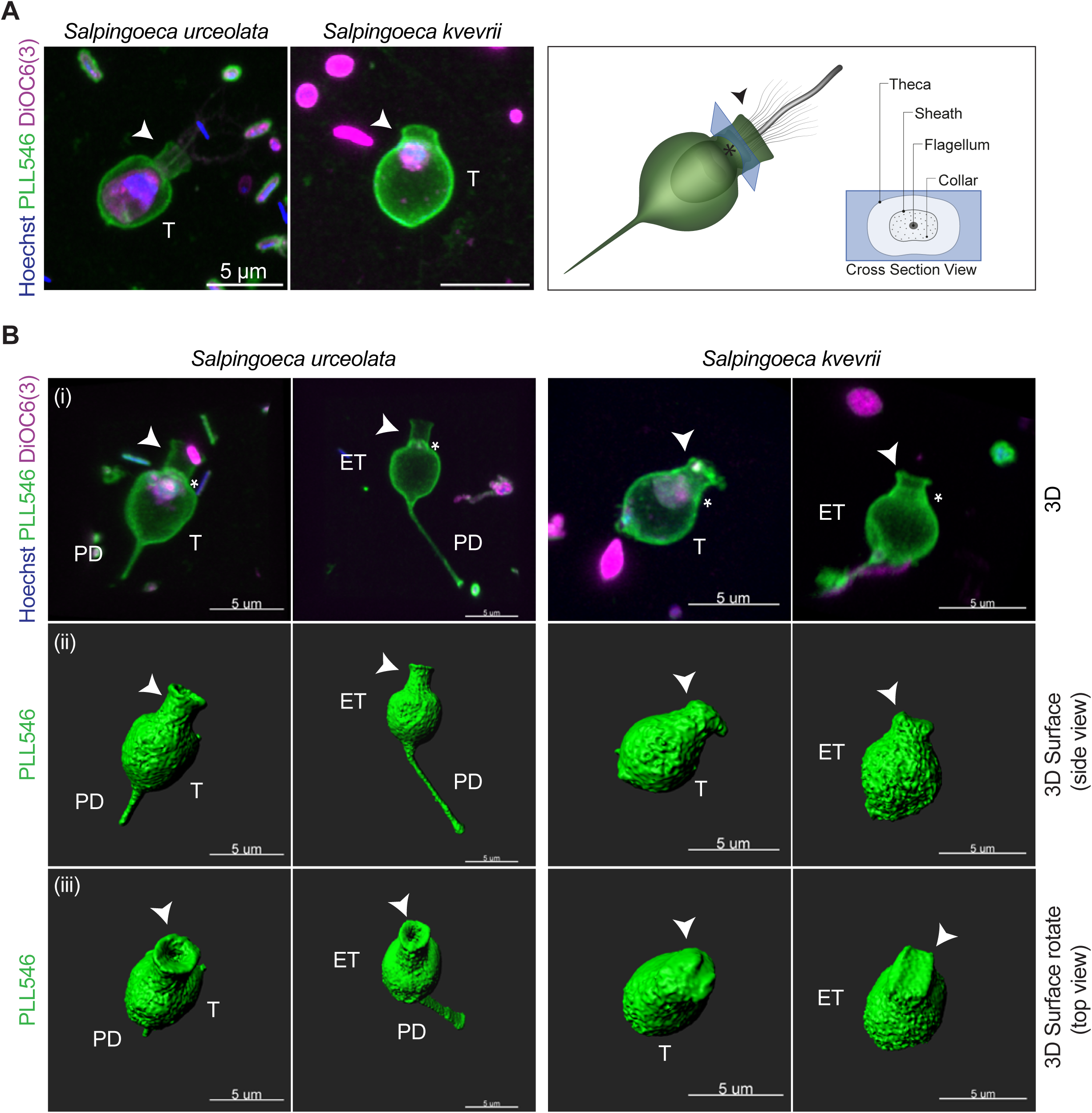
Comparative analysis of theca morphology in *Salpingoeca urceolata* and *Salpingoeca kvevrii* using super-resolution fluorescence microscopy stained with Hoechst (blue, DNA), PLL546 (green, extracellular structures), and DiOC6(3) (magenta, cell membranes). Cells are co-cultured with their co-isolated mixed bacterial prey. T: theca, ET: empty theca, PD: pedicel. Scale bars: 5 μm. **A)** Super-resolution images of individual cells. Both species exhibit a theca. White arrowheads indicate the distinct neck of the theca. **B)** 3D reconstructions of thecate cells and empty thecae from each species. Three visualizations are shown: (i) Overlay of all three fluorescence channels, (ii) 3D surface reconstruction of PLL546 staining, and (iii) Rotated view of panel (ii). White arrowheads indicate the distinct neck of the theca. Asterisks mark a putative inner structure sheath-like layer. Both *S. urceolata* and *S. kvevrii* display a flask-shaped theca with species-specific differences in neck length, and the presence of a pedicel in *S. urceolata* only.

An additional inner sheath structure has been reported by electron microscopy in flask-shaped thecate choanoflagellates including *S. urceolata*, potentially serving to anchor the theca to the cell body (Carr et al., 2008; Leadbeater, 1977). In our study, the accumulation of staining at the base of the neck in both *S. kvevrii* and *S. urceolata* (**Figure 5B**, indicated by an asterisk) may correspond to this structure, although further ultrastructural confirmation is needed.

### 5. The tubular-shaped theca of *Daedalina diplosolina*, with a twist

A novel choanoflagellate genus and species, *Daedalina diplosolina*, was isolated from a freshwater mangrove in the Caribbean island of Grenada. Phylogenomic analysis confirmed that the species clusters within Clade 3 with strong support, as a sister to the group/clade containing *Salpingoeca dolichothecata* and *S. tuba* (**Figure 1A** and **Supplementary Figure 1**). The tubular-shaped theca of *D. diplosolina* observed during DIC experiments (**Figure 1B**) matched that of other Clade 3 members, corroborating its phylogenetic affiliation. Subsequent super-resolution microscopy and 3D reconstruction with PLL546 confirmed the tubular shape of the extracellular covering in *D. diplosolina* (**Figure 6A-B**). However, SEM observation revealed an unexpected structural feature: the theca consists of two distinct parts, joined along a linear connection (**Figure 6C** and **Supplementary Figure 4A-i-ii**). To ensure this was not a fixation artifact, the presence of this junction was also confirmed in live cells (**Supplementary Figure 4B-i**).

**Figure 6:**
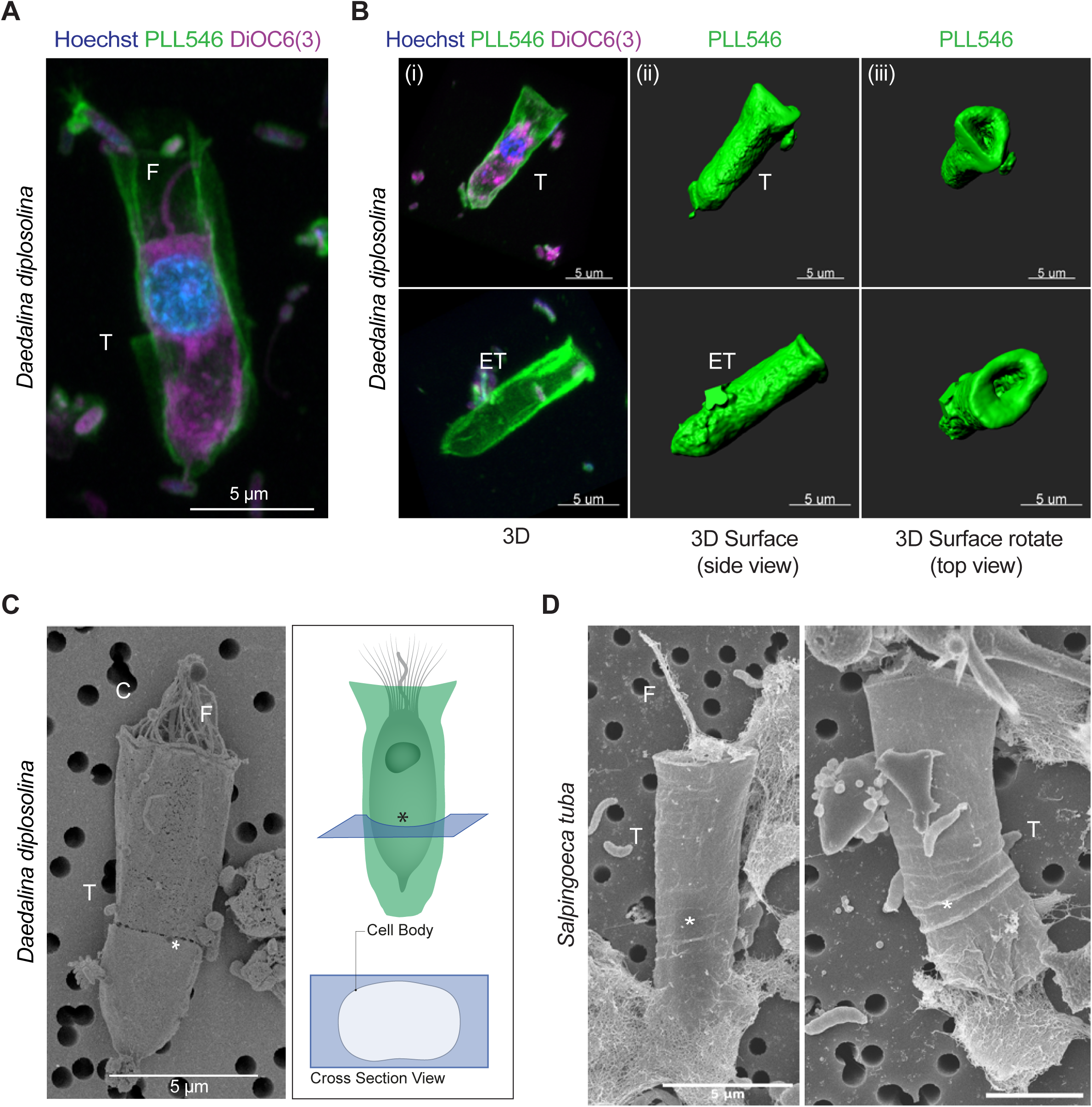
***Daedalina diplosolina*, a Clade 3 choanoflagellate with a two-part tubular theca and a comparison to *Salpingoeca tuba*, also from Clade 3. with co-cultured bacterial prey. Hoechst (blue): DNA content, PLL546 (green): extracellular structures, DiOC6(3) (magenta): membranes. T: theca, F: flagellum, C: collar. ET: empty theca. Scale bar: 5 μm.** **A)** Super-resolution microscopy of individual thecate *D. diplosolina* cell. **B)** 3D reconstructions of thecate cell and empty thecae. Three visualizations are shown: (i) Overlay of all three fluorescence channels, (ii) 3D surface reconstruction of the PLL546 staining, and (iii) Rotated view of panel (ii). **C)** Scanning electron microscopy (SEM) image and drawing of the theca of *D. diplosolina*. The tubular theca is composed of two parts. An asterisk indicates the junction of the two parts. **D)** Scanning electron microscopy (SEM) images of the theca of *S. tuba.* The asterisk indicates an accordion-like flexible region.

To determine whether this morphology was characteristic of Clade 3 more broadly, *Salpingoeca tuba* was re-isolated (strain BEAP0445) and examined using SEM (**Figure 6D)**. In contrast to *D. diplosolina*, no evidence of a distinct junction or bipartite organization was observed. Instead, the extracellular structure of *S. tuba* contained an accordion-like flexible region (labelled with an asterisk) capable of deformation while maintaining structural continuity. These observations suggest that the conspicuous separation observed in *D. diplosolina* is not a general feature of Clade 3. Nevertheless, both species possess modifications that may increase flexibility within an otherwise relatively long tubular extracellular architecture, although they appear to achieve this through different structural mechanisms.

SEM analysis revealed additional morphological variation in *D. diplosolina*. Alongside typical thecate cells with an intact two-part theca, cells were observed within only the upper part (**Supplementary Figure 4A-iii**), or within intact thecae that extended into pedicels (**Supplementary Figure 4A-i**).

Live cell imaging confirmed these morphotypes and indicated further diversity within the culture (**Supplementary Figure 4B**). Because cells are strongly adhered to the substrate, mechanical scraping was applied before imaging. Following this treatment, a variety of morphotypes were observed, including intact thecate cells (**Supplementary Figure 4B-i**), empty thecae (intact, upper or lower, **Supplementary Figure 4B-ii-v-vi**), cells with only the upper thecal part (**Supplementary Figure 4B-iv**) and naked cells, some of which were free swimming in the water column (**Supplementary Figure 4B-vii**) while others were attached to the surface with filopodia (data not shown). Finally, crawling cells, characterized by a short collar, a long flagellum and filopodia, were also present (**Supplementary Figure 4B-viii**).

After one week in culture following manual scraping, *D. diplosolina* populations comprised intact thecate cells, naked cells and cells retaining only the upper part of the theca, although their relative proportions differed between the two experiments performed under different culture conditions (**Supplementary Figure 4C**). Notably, re-applying mechanical disruption appeared to increase the proportion of naked cells, particularly those missing the lower part of the theca. These initial observations raise the possibility that the two-part theca may facilitate partial shedding (fast escape of the cell) in response to mechanical stress, although this remains to be tested. This interpretation is consistent with the environmental origin of the culture, as mangrove environments in general are characterized by strong fluctuations in salinity, oxygen availability and hydrodynamic disturbance (Alongi, 2009; Wolanski, 1992).

Further observation of the crawling cells (**Supplementary Figure 4B-viii**), revealed that they moved in the direction opposite to the flagellum, while the flagellum itself appeared immotile. Super-resolution microscopy confirmed the presence of filopodia and highlighted the distinct features of the crawling cells compared to thecate cells (**Figure 7A**): a smaller-sized cell body, longer flagellum and a shorter collar of microvilli.

**Figure 7:**
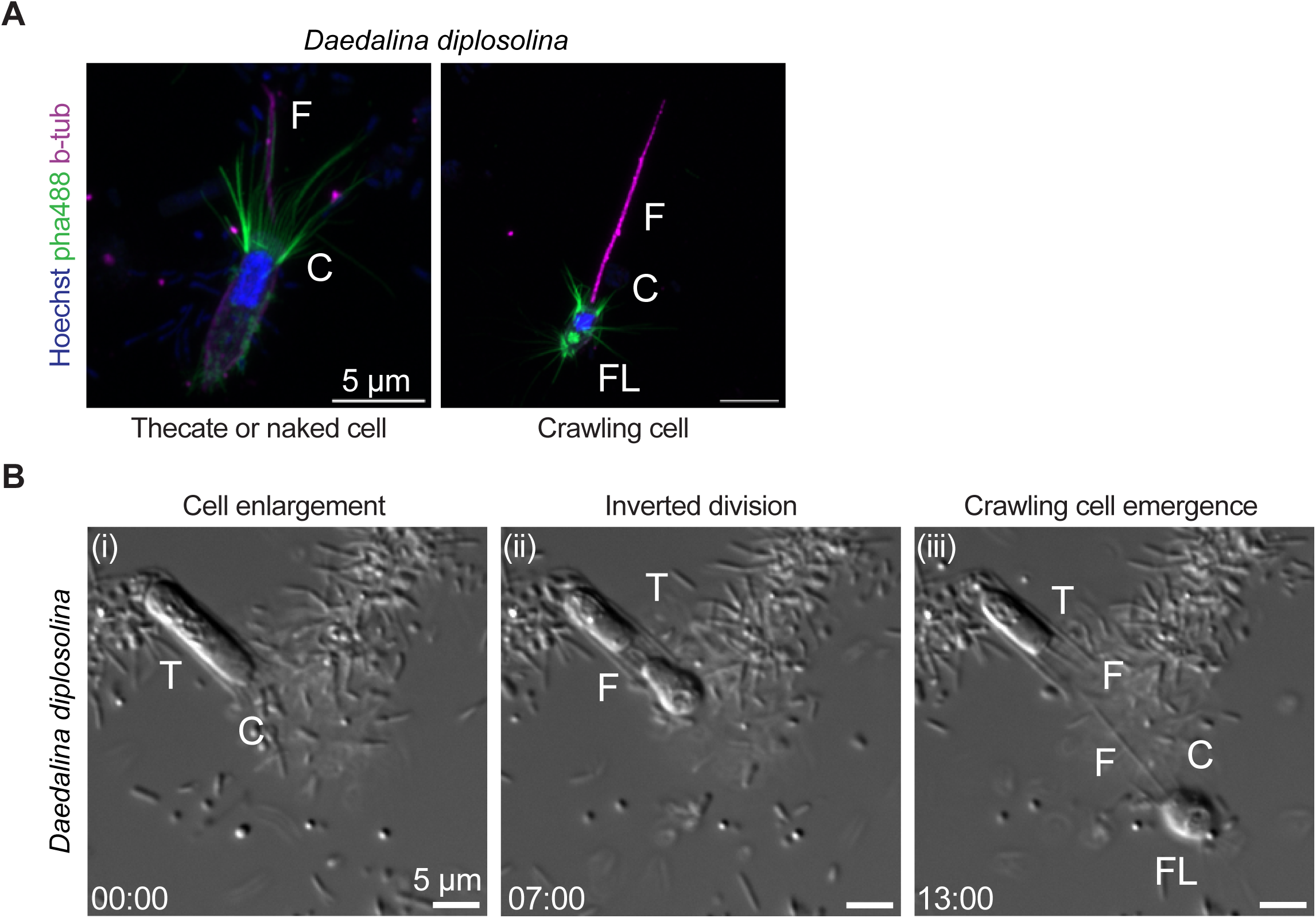
Origin of the crawling cell phenotype in *Daedalina diplosolina*. T: theca, C: collar, F: flagellum, FL: filopodia. **A)** Super-resolution microscopy of DNA (Hoechst, blue), actin structures (phalloidin, green), and flagella (β-tubulin, magenta) highlights differences between thecate or naked cells and crawling cells, the latter characterized by a smaller cell body, longer flagellum, shorter collar, and filopodia. Scale bars: 5 μm. **B)** Live-cell imaging sequence shows that crawling cells emerge via division of thecate cells: cell enlargement (i), inverted division with daughter cells positioned flagellar pole to flagellar pole (ii), and emergence of a crawling daughter cell while the other remains thecate (iii). Three independent division events of this type were captured via time-lapse imaging, although this division pattern of thecate cells was observed consistently during culture maintenance. Time is indicated in MM:SS.

Live imaging further revealed that the crawling cells emerged from the division of thecate cells (**Figure 7B**). More specifically, before division, the cell within the theca expanded in size (**Figure 7B-i**), followed by division in an inverted pattern (**Figure 7B-ii**), with the daughter cell flagellar poles facing each other during separation. In the end of the division, the inward daughter remains thecate, while the outward exits the theca in the crawling form (**Figure 7B-iii**).

Interestingly, cell division in choanoflagellates has been thought to be largely influenced by the type of extracellular covering. In craspedids, two division modes have been observed: (1) species with non-constraining coverings divide vertically (longitudinal division), while (2) thecate species typically undergo emergent division, during which they become amoeboid and partially emerge from the parent theca and then divide longitudinally (Leadbeater, 2014). In contrast, *D. diplosolina* represents a case of inverted division in a thecate species. After division, crawling cells actively explore their environment before settling and producing a new theca (see video in **Supplementary material 15**). Interestingly, crawling cells were also observed in *S. tuba*, suggesting that this behaviour may be shared more broadly among members of Clade 3.

### 6. *Hippolytella eleftheri* offers insights into the timing of the evolution of the lorica

*Hippolytella eleftheri* was isolated from the banks of the Fraser River in Vancouver, BC and represents one of the first cultured representatives of the freshwater associated group of choanoflagellates, the FRESCHO–3 clade. This group was initially identified from environmental 18S rRNA gene surveys and was shown to form a putative sister group to the Acanthoecida, one of the two main choanoflagellate orders (**Figure 8A**) (Del Campo and Ruiz-Trillo, 2013).

**Figure 8:**
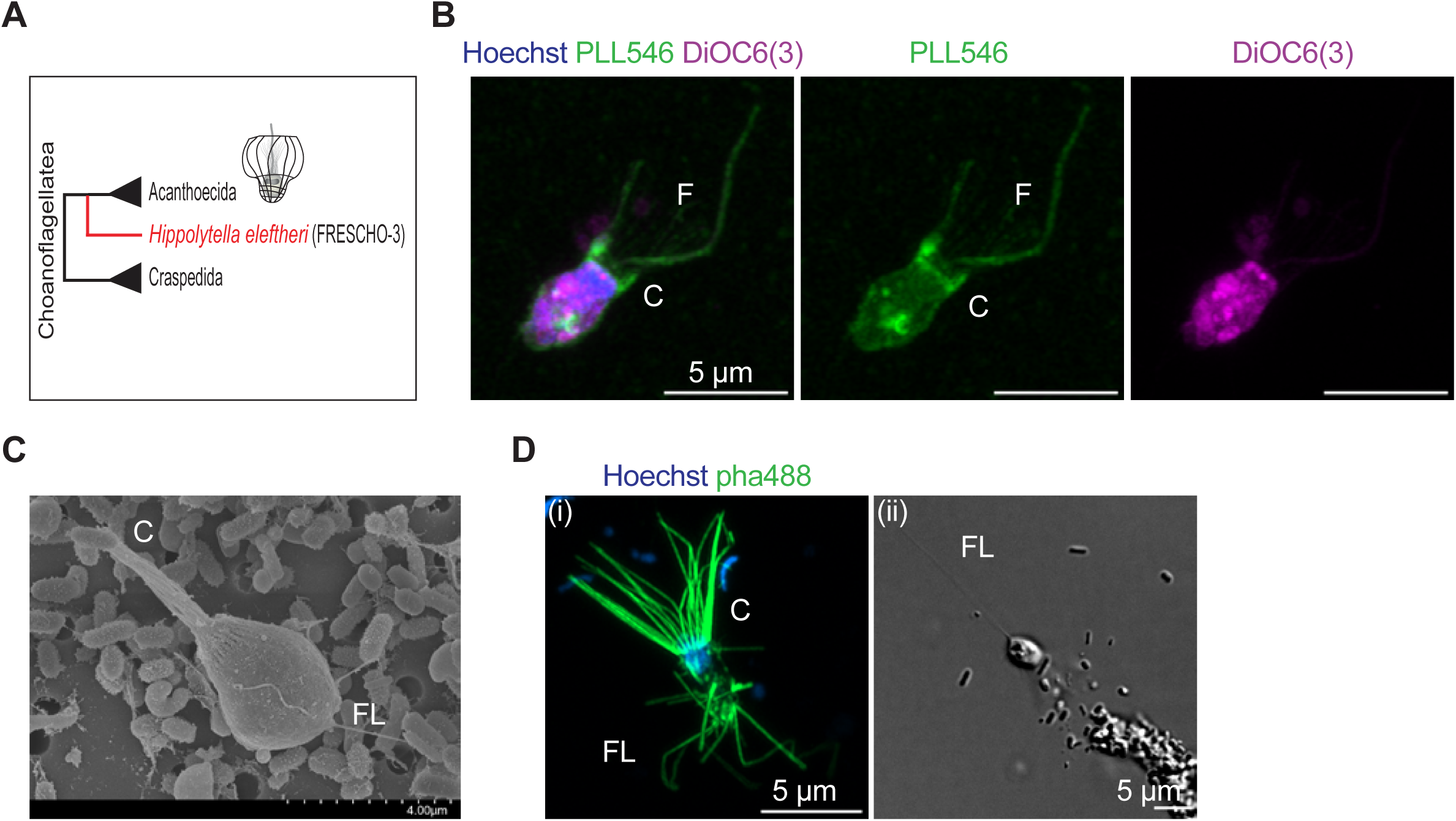
***Hippolytella eleftheri*, a representative of the FRESCHO-3 group (a sister group to Acanthoecida), does not possess a lorica. F: flagellum, FL: filopodia, C: collar. *H. eleftheri* is co-cultured with its bacterial prey.** **A)** Illustration showing the phylogenetic position of *H. eleftheri* and the FRESCHO-3 group, as sister to all loricate species. **B)** Super-resolution microscopy of individual *H. eleftheri* cell stained with Hoechst (blue): DNA content, PLL546 (green): extracellular structures, DiOC6(3) (magenta): membranes. The PLL546 staining shows a diffuse signal closely following the cell membrane, suggesting the absence of an extracellular covering. Scale bar: 5 μm. **C)** Scanning electron microscopy (SEM) image of *H. eleftheri* confirming the absence of the lorica and the presence of filopodia. Scale bar: 4 μm. **D)** Super-resolution microscopy (i) and DIC microscopy (ii) of an individual *H. eleftheri* cell stained with Hoechst (blue): DNA content, pha488: actin-based structures. All cells examined displayed filopodia (n = 30), which could extend from the entire cell body and reach considerable length. Scale bar: 5 μm.

Transcriptome generation and phylogenetic analysis confirmed, with strong support, that *H. eleftheri* was positioned as a sister group to the rest of the Acanthoecida (**Figure 1A**). Subsequent phylogenetic analysis incorporating the 18S and 28S rRNA gene sequences of another representative of the FRESCHO-3 group, choanoflagellate sp. strain FN-2015 (Jeuck, 2014) showed that it is sister to *H. eleftheri* (**Supplementary Figure 1**).

The absence of a lorica in *H. eleftheri* caught our attention, as the cells appeared naked and lacked any visible extracellular covering (**Figure 1B**). Consistent with the DIC observations, we did not detect a lorica by fluorescence microscopy. Instead, PLL546 staining produced a diffuse signal that closely outlined the cell membrane and extended only slightly beyond the microvilli (**Figure 8B**). This pattern most likely reflects direct interaction of PLL546 with the cell surface, but it could also be consistent with a very thin, tightly fitting sheath-like covering or another nonsiliceous type of extracellular material. As a positive control, PLL546 staining clearly labelled the lorica of the acanthoecid *Acanthocorbis camarensis* (**Supplementary Figure 5**).

The absence of a lorica was further confirmed by SEM, which revealed thin filopodia extending from the cell surface (**Figure 8C**). To verify the presence of these filopodia, we stained cells with phalloidin, which labels F-actin rich structures such as filopodia and the collar, together with Hoechst for DNA staining (**Figure 8D-i**). This confirmed that the observed projections correspond to actin-rich filopodia, extending across the entire cell surface. Live imaging showed that some cells use these filopodia for movement, sometimes extending particularly long filopodial projections up to 4 times the length of the cell body (**Figure 8D-ii**). We also observed swimming cells retaining filopodia, suggesting a broader role for these structures across different behavioural states (data not shown).

Previous work on FN-2015 reported the absence of a lorica, even when the species was cultured in different media designed to promote lorica formation (Jeuck, 2014). Consistent with these observations, in our experiments, *H. eleftheri* was grown in freshwater RS medium (recipes can be found in MediaDive: <u>P4</u>, <u>P5</u>), a medium whose components, including silica, support lorica formation in other loricate choanoflagellates (Gàlvez-Morante et al., 2026) and silica-based extracellular structure production in diatoms (unpublished observations from laboratory cultures), yet no lorica was observed. These findings suggest that the absence of a lorica is a common feature of the FRESCHO–3 group, rather than a species-specific adaptation.

Transcriptome analysis of *H. eleftheri* revealed the absence of silicon transporter genes, which are thought to be required for lorica formation (Li et al., 2018; Marron et al., 2016); silicon transporters are among the most highly expressed genes in other loricate choanoflagellates (Marron et al., 2016; Richter et al., 2018), further supporting that *H. eleftheri* lacks the molecular capacity to produce a lorica. Additionally, silicon transporter genes are also absent from the single-cell amplified genome (SAG) of UC4 (Uncultured Choanoflagellate species 4), which is thought to branch as the earliest-diverging sister lineage to Acanthoecida (López-Escardó et al., 2019).

## Discussion

### The evolution of extracellular structures in choanoflagellates

The comparative analysis presented in this study shows that, unsurprisingly, extracellular structures are often conserved among closely related choanoflagellate species. In contrast, broader morphological divergence becomes evident when comparing more distantly related clades. Historically, the terminology used to describe choanoflagellate extracellular coverings has been inconsistent, largely because the biochemical composition of most structures remains poorly characterized. Following previous efforts to simplify the terminology (Leadbeater, 2014), we broadly distinguish between flexible extracellular structures, including glycocalyces and sheath-like coverings, as opposed to rigid extracellular structures, including organic thecae and siliceous loricae. While these categories differ in morphology and composition, they should not necessarily be viewed as entirely independent extracellular innovations. Instead, our observations suggest that choanoflagellate extracellular morphologies may represent a continuum, ranging from flexible coverings closely associated with the cell surface to highly structured rigid architectures. Future efforts aimed at characterizing the biochemical composition of these structures will be critical not only for refining our understanding and enabling more targeted staining approaches, but also for assessing whether extracellular structures that appear morphologically similar across choanoflagellates and Metazoa share common molecular foundations. Resolving these biochemical similarities and differences will be essential for distinguishing homologous extracellular systems inherited from a common ancestor from analogous structures that evolved convergently in response to similar selective pressures.

Consistent with this interpretation, flexible extracellular coverings appear to be widespread across choanoflagellates, occurring not only in non-thecate craspedids (such as the *Hartaetosiga* and *Codosiga* genera) but also in thecate species (such as *Choanoeca flexa, Salpingoeca kylix, Salpingoeca rosetta* and *Oinochoeca dipylis*) and even in some loricate acanthoecids, including *Parvicorbicula quadricostata* and *Calliacantha natans* (Dayel and King, 2014; Hibberd, 1975; Leadbeater, 2014, 1977, 1973, 1972; Leadbeater and Morton, 1974). Similar sheath-like structures have also been reported in species assigned to the craspedid genera *Diploeca* and *Diplosiga*, although the species have never been isolated and these observations are based primarily on historical light microscopy studies (Ellis, 1930; Frenzel, 1891). The repeated occurrence of these flexible coverings across phylogenetically distant lineages suggests that they may represent an ancient feature of choanoflagellate biology (**Figure 9**). Their existence both independently and in association with rigid extracellular structures suggests that their functions may vary among species, potentially contributing to cell protection, environmental interactions, feeding, hydrodynamics, attachment and the association between the cell body and surrounding extracellular material. In Metazoa, the sister group to choanoflagellates, glycocalyx layers are present on many cell surfaces possessing microvilli (**Figure 9**), including brush border cells, choanocytes and sensory hair cells, although they are not described on all microvillar cells (Mah et al., 2014; Sharkova et al., 2023; Sun et al., 2020). Cells lacking organized microvilli, such as mesenchymal cells, can also be surrounded by a glycocalyx-like pericellular coat (Evanko et al., 2007). These layers could potentially differ in molecular composition and biological context, but they share the common feature of forming flexible extracellular interfaces positioned at the boundary between cells and their environment. Within Choanozoa (the lineage comprising animals and choanoflagellates), the occurrence of glycocalyx-like structures in choanoflagellates together with the presence of glycocalyx layers on microvilli-bearing metazoan cell types (although the morphology and function of collar cells diversified substantially in Metazoa) is consistent with the hypothesis that the choanozoan ancestor possessed a flexible extracellular covering associated with the collar complex.

**Figure 9:**
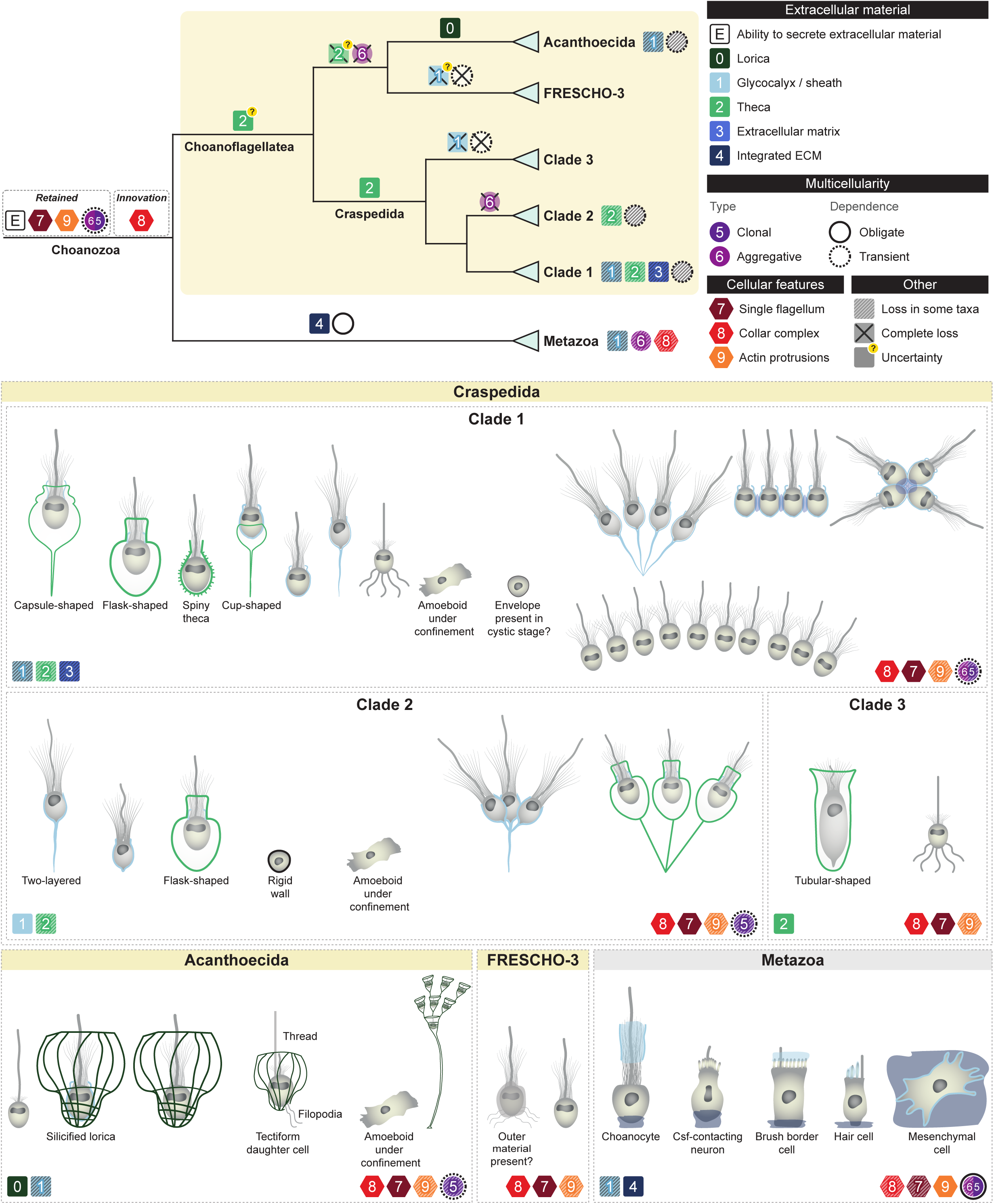
Evolution of extracellular structures, cellular traits and multicellularity in Choanozoa. The phylogeny (top left) summarizes the relationships between Metazoa and the major choanoflagellate lineages (yellow background) and the distribution of extracellular structures, cellular traits and multicellularity across the group. Traits inferred for the choanozoan ancestor are divided into those retained from earlier opisthokont ancestors versus the innovation of Choanozoa, the collar complex. Representative cell morphologies (bottom; not to scale) illustrate the principal extracellular architectures of each lineage. Squares represent extracellular structures, with the open square (E) indicating the capability to secrete extracellular material. Because the composition of most holozoan extracellular structures is currently unknown, a single symbol may represent different structures with similar morphology and does not necessarily imply homology. Hexagons show conserved cellular features, filled circles denote the type of multicellular organization (clonal or aggregative), and circle outlines indicate multicellular dependency (obligate or transient). Hatched symbols indicate partial losses in descendant taxa, crosses (×) indicate inferred complete losses, and question marks indicate uncertainty.

In contrast to the widespread distribution of flexible extracellular coverings, our data suggest that siliceous loricae represent a derived extracellular innovation that evolved within the acanthoecid lineage (**Figure 9**), based on the absence of both a lorica and silicon transporter genes in acanthoecid sister species *Hippolytella eleftheri*, the absence of a lorica in FN-2015 and the lack of these genes in the SAG of UC4.

Pinpointing the evolution of the theca is less straightforward (**Figure 9**). Unlike loricae, whose evolutionary origin can be linked to the presence and duplication of silicon transporter genes to produce sufficient intracellular concentrations for siliceous biomineralization, the composition of most choanoflagellate organic extracellular structures remains poorly understood. Consequently, it is difficult to determine whether thecae, sheaths and glycocalyces represent distinct extracellular architectures or different structural organizations of the same underlying extracellular materials. Nevertheless, the apparent restriction of rigid organic extracellular structures to craspedid choanoflagellates suggests that they may represent a derived feature of this lineage. At the same time, their absence in genera such as *Hartaetosiga*, *Codosiga*, *Monosiga* and *Mylnosiga* indicates that these structures have likely been lost repeatedly during craspedid evolution, although it should be noted that *Codosiga*, which lacks a theca, retains the ability to produce a rigid extracellular enclosure during cyst formation (Stoupin et al., 2012). Similar cyst formation involving the deposition of an extracellular wall has also been reported in a freshwater choanoflagellate, *Desmarella moniliformis* (Leadbeater and Karpov, 2000), and in *Choanoeca flexa* following desiccation, although no associated extracellular structures have been identified (Ros-Rocher et al., 2026).

Despite these repeated losses within the craspedid lineage, the overall morphology of thecae often appears remarkably conserved among closely related species. For example, *Salpingoeca kylix* and *S. rosetta* both possess cup-shaped thecae (**Figure 3**), while *Oinochoeca dipylis* and *Stagondoeca pyriformis* share characteristic capsule-shaped morphologies (**Figure 4** and Supplementary **Figure 3**). Similarly, *Salpingoeca urceolata* and *S. kvevrii* exhibit matching flask-shaped structures despite differences in pedicel morphology (**Figure 5**). Within Clade 3, the newly isolated strain *Daedalina diplosolina* possesses a tubular theca structure (**Figure 6**) similar to those reported in other representatives of the clade, suggesting that this morphology may also be evolutionarily conserved.

At broader evolutionary scales, however, considerable diversification becomes apparent (**Figure 9**). Clade 3 appears to be dominated by tubular morphologies, whereas Clade 2 is characterized primarily by flask-shaped forms. In contrast, Clade 1 exhibits a broader range of extracellular architectures, including cup-shaped, capsule-shaped and flask-shaped thecae, as well as the spiny theca of *Lagenoeca antarctica* (Nitsche et al., 2007). The occurrence of flask-shaped thecae in several Clade 1 species, including *Choanoeca flexa* and *C. perplexa*, together with their predominance within Clade 2, suggests that this morphology either evolved repeatedly or was already present in the common ancestor of Clades 1 and 2 before undergoing subsequent modification in descendant lineages.

Taken together, these observations support the hypothesis that the last common choanoflagellate ancestor possessed an extracellular covering, likely resembling a sheath or glycocalyx-like structure. The widespread distribution of such coverings across phylogenetically distant lineages, including species with and without rigid extracellular architectures, is consistent with an ancestral origin. Whether this ancestor could also organize extracellular material into a rigid theca-like structure remains uncertain, particularly given our limited knowledge of the composition and homology of organic coverings. More elaborate extracellular morphologies, including siliceous loricae, appear to represent lineage-specific modifications and specializations of this ancestral extracellular system.

### Extracellular structures influence cell division and colony formation

Beyond their structural and protective roles, as well as the recently proposed role of the thecate stage in nutrient uptake in *Salpingoeca rosetta* (Leon et al., 2025), extracellular coverings in choanoflagellates may also have played an important role in the evolution of multicellular life history stages (**Figure 9**). In several species, extracellular material appears to be closely associated with multicellular organization, either through direct physical attachment mediated by extracellular structures such as loricae (*Crinolina aperta*), collars (*Choanoeca flexa*), theca- (*Salpingoeca urceolata*) or glycocalyx- derived pedicels (the *Hartaetosiga* genus) or through the secretion of additional extracellular matrix components (Brunet et al., 2019; Dayel et al., 2011; Escalera et al., 2019; Leadbeater and Morton, 1974; Levin et al., 2014; Richter et al., 2018). In *S. rosetta*, for example, rosette colonies require the secretion of Rosetteless, a C-type lectin-like extracellular matrix protein. Extracellular material also contributes to multicellular organization in other holozoans. In filasterea, *Capsaspora owczarzaki* extracellular material secreted during aggregative stages contributes to cell adhesion and multicellular aggregation (Sebé-Pedrós et al., 2013b), and a gene with homology to collagen IV is upregulated in *Ministeria vibrans* aggregative colonies (Li et al., 2026). Integrated extracellular matrix systems diversified in Metazoa, composed primarily of collagens, laminins, fibronectins and proteoglycans (Fidler et al., 2017; Hynes, 2012, 2009; Özbek et al., 2010). Unlike the localized coverings and aggregation-associated extracellular materials observed in other opisthokonts, metazoan ECM forms continuous tissue-level scaffolds that coordinate interactions among multiple cell types and play central roles in morphogenesis, cell differentiation and tissue organization.

Our observations of *Hartaetosiga kampana* further support a close relationship between extracellular structures and multicellular organization. More specifically, cells of *H. kampana* are not homogeneous, but instead occur in distinct life stages, including stalk-associated colonies. This ability to form colonies through cell division has previously been reported within the *Hartaetosiga genus* (Leadbeater and Morton, 1974), although the cues responsible for its induction remain unknown. Current molecular understanding of colony formation in choanoflagellates largely derives from a limited number of model organisms, particularly *Salpingoeca rosetta* and *Choanoeca flexa* (Alegado et al., 2012; Brunet et al., 2019; Fairclough et al., 2010; Reyes-Rivera et al., 2022; Ros-Rocher et al., 2026). However, none of these established models form colonies attached through stalks, a mode of organization observed not only in the *Hartaetosiga* genus within Clade 1, but also in the *Codosiga* genus within Clade 2, as well as in some acanthoecid species such as *Polyoeca dichotoma* (Boucaud-Camou, 1966; Leadbeater and Morton, 1974; Stoupin et al., 2012). The repeated observation of stalk-associated colonies in both craspedids and acanthoecids further raises the possibility that this form of multicellular organization represents a recurrently favored architectural strategy within choanoflagellates.

Another aspect of choanoflagellate biology that appears to be strongly influenced by the nature of the extracellular covering is the mode of division (Leadbeater, 2014). Species lacking rigid extracellular structures typically divide longitudinally (Karpov and Leadbeater, 1997), whereas rigid extracellular architectures preclude classical longitudinal division and instead require alternative strategies to facilitate daughter-cell release. These include longitudinal division following theca exit by adopting an amoeboid morphology in many craspedids, diagonal division in nudiform acanthoecids and inverted division in tectiform acanthoecids (Leadbeater, 2010, 1994a, 1977; Leadbeater et al., 2008). Division mode itself can be plastic: the tectiform *Stephanoeca diplocostata* shifts from inverted to nudiform-like division following silicon depletion (Leadbeater and Carr, 2022), indicating that division mode can respond to changes in extracellular architecture within a single lineage.

This interpretation is further supported by the FRESCHO-3 representative *Hippolytella eleftheri*. Although this lineage is the sister group to acanthoecids, it lacks a lorica and appears to undergo a more conventional longitudinal division. In contrast, the craspedid *Daedalina diplosolina* appears to exhibit a division pattern resembling the inverted division observed in tectiform acanthoecids despite belonging to the distantly related Clade 3. The opposing orientation of the daughter cells during cytokinesis and the subsequent emergence of a crawling stage suggest that similar structural constraints may have repeatedly shaped the evolution of division behaviour across choanoflagellates. The long flagellum observed in *D. diplosolina* may additionally facilitate daughter-cell detachment, potentially fulfilling a role analogous to the structural thread described in some loricate species (Leadbeater, 1994a, 1979).

If the last common choanoflagellate ancestor indeed possessed a flexible extracellular covering, as proposed above, then longitudinal division likely represents the ancestral mode of cell division, with alternative division strategies evolving later, potentially via reuse of the same retained genes or pathways, in association with increasingly rigid extracellular architectures. Such a flexible covering may also have facilitated the retention of daughter cells following division, providing a simple mechanism for the formation of multicellular colonies. Nevertheless, colony formation can also occur in species with rigid extracellular coverings, indicating that a flexible covering is not a prerequisite for multicellularity (Escalera et al., 2019; McKenzie et al., 1997; Richter et al., 2018).

Outside Choanozoa, extracellular structures are morphologically diverse across Opisthokonta, suggesting that the capacity to secrete extracellular material appears to have been retained in this group (**Figure 9**). Within Holomycota, fungi are characterized by the presence of cell walls containing chitin and glucans that provide structural support and protection while facilitating multicellular growth and the osmotrophic lifestyle of the group (Gow et al., 2017; Gow and Lenardon, 2023). These walls are frequently associated with outer fibrillar layers rich in glycoproteins and mannoproteins. However, cell wall composition varies among fungi and many fungal zoospores lack a wall (for example those of chytrids), showing that rigid coverings can also be restricted to particular life stages within this group (Richards et al., 2017). Aphelids and Rozellids, two paraphyletic lineages sister to fungi, also form extracellular walls during specific life history stages (Karpov et al., 2014; Mikhailov et al., 2022; Torruella et al., 2018). Their genomes encode conserved components of cell wall synthesis and remodelling, and chitin has been detected in the walls of the invasive cysts. The cyst wall has been proposed to be essential for host invasion, providing a rigid support from which the infection tube penetrates the host cell wall, after which the parasitoid develops as a wall-less plasmodium within the host. Many nucleariids possess a mucous extracellular coat (sometimes labeled as glycocalyx), which can vary with environmental conditions and harbour bacteria ectosymbionts (Gabaldón et al., 2022). In some genera, this coat also incorporates additional extracellular particles, including endogenous siliceous scales in *Pompholyxophrys* and exogenous material in *Lithocolla*. In addition, most nucleariids can form resting cysts surrounded by one or more thick layers of extracellular material, possibly derived from the mucous coat.

Within Holozoa, ichthyosporeans include walled coenocytic stages, within which repeated nuclear divisions and cellularization occur prior to the release of daughter cells, leaving the extracellular enclosure behind (Dudin et al., 2019; Marshall and Berbee, 2011; Shabardina et al., 2024). These developmental stages are associated with a largely absorptive or osmotrophic lifestyle and the retention of a diverse repertoire of chitin synthases together with evidence of chitin staining in their walls further suggests functional and evolutionary similarities with fungal extracellular compartmentalization (Torruella et al., 2015). *Tunicaraptor* displays a close-fitting extracellular envelope (referred to as a theca) decorated with filamentous projections (Tikhonenkov et al., 2020b), whereas pluriformeans possess extracellular coverings associated with particular life history stages. For example, scale-like extracellular structures observed by electron microscopy in the coenocytic stage of *Corallochytrium limacisporum* (Raghu-kumar, 1987) and outer envelopes enclosing spherical cysts of *Syssomonas multiformis*, within which successive cell divisions can produce many naked flagellated daughter cells (Tikhonenkov et al., 2020a).

### Actin based structures and behavioural plasticity

Another important behavioural and cytoskeletal feature highlighted by our observations is the widespread occurrence of filopodia across choanoflagellates, which suggests that the last common choanoflagellate ancestor likely was capable of both free swimming and substrate-associated locomotion.

More specifically, in *Daedalina diplosolina*, daughter cells emerging from inverted division displayed prominent filopodia which enabled crawling. Occasional crawling cells were also observed in *Oinochoeca dipylis* when actively exploring the substrate **(Supplementary Figure 3B**), while in the FRESCHO-3 strain *Hippolytella eleftheri*, filopodia were present in most observed cells, including both attached and swimming stages. Cells extending filopodia have also been described in several distantly-related choanoflagellates. For example, in *Salpingoeca rosetta*, fast-swimming cells use filopodia both to attach to empty thecae, which serve as scaffolds for the formation of new thecae, and to facilitate detachment during theca exit (Dayel et al., 2011). Filopodia have also been observed during *S. rosetta* colony formation, where they may contribute to cell-cell interactions and colony organization. Similarly, the recently described hyperhalophilic species *“Salpingoeca” surira, “Salpingoeca” huasca and “Salpingoeca” prava* all display life history stages characterized by prominent filopodia (these species were described with their generic names in quotes to reflect the fact that the genus *Salpingoeca* is in need of revision; (Schiwitza et al., 2018)). Earlier studies also described filopodia-bearing species such as *Kentrosiga echinobasis*, which can form chain-like colonies while retaining these protrusions (Tong, 1997). In addition, crawling daughter cell stages are well documented in tectiform acanthoecids such as

*Stephanoeca diplocostata* and *Diaphanoeca grandis* (Leadbeater, 1994b; Leadbeater and Cheng, 2010), as well as in craspedids such as *Monosiga ovata* and *Hartaetosiga gracilis* (Leadbeater, 2014; Leadbeater and Morton, 1974). Furthermore, under physical confinement, choanoflagellates can adopt amoeboid morphologies, revealing an additional degree of cellular plasticity (Brunet et al., 2021).

Interestingly, while actin-based protrusions show considerable diversity in morphology and function, the actin-supported collar of microvilli remains remarkably conserved across choanoflagellates. Actin-based protrusions are also widespread throughout other holozoan lineages (Sebé-Pedrós et al., 2013a), and their presence in Asgard archaea suggests a deeper evolutionary origin, potentially predating the last eukaryotic common ancestor (LECA) (Rodrigues-Oliveira et al., 2023). These structures likely formed part of the ancestral cellular toolkit inherited by both choanoflagellates and animals. From this perspective, the choanoflagellate collar can be viewed as an evolutionary innovation, transforming a general actin-based protrusion system into a highly organized feeding apparatus that became a defining feature of the group and may have contributed significantly to its ecological success and diversification.

## Materials and Methods

### Isolation and maintenance of the choanoflagellate strains

*Hartaetosiga kampana* (BEAP0001, CRO2OCHO1) was isolated in July, 2014 from shallow marine sediment collected in the Malo Jezero lagoon, Mljet Island, Croatia (Adriatic Sea, GPS: 42.784916, 17.350350). The sample was taken from the top 10 cm of aerobic sediment and stored at −20 °C until February 2020. Enrichment was performed in 0.1% YT medium (100 mg yeast extract, 200 mg tryptone in 100 ml distilled water) prepared in sterile natural seawater and incubated at room temperature.

*Oinochoeca dipylis* (BEAP0002, LJO2) was collected in August, 2020 from Playa del Janubio, Lanzarote, Canary Islands, Spain (Atlantic Ocean, GPS: 28.935796, −13.830647), from the top 10 cm of aerobic sandy sediment. Enrichment was also conducted in 0.1% YT medium in sterile natural seawater at room temperature.

For both *Hartaetosiga kampana* and *Oinochoeca dipylis*, single choanoflagellate cells were manually isolated using micromanipulation and transferred to 24-well plates containing sterile seawater supplemented with 0.1% YT medium. Cultures were initially maintained at 15 °C and later transitioned to marine RS medium (1:100 P5:P4, recipes can be found in MediaDive: <u>P4</u>, <u>P5</u>) at 17 °C in darkness. To limit the diversity and density of bacterial prey, each culture was treated with a panel of different antibiotics to promote optimal choanoflagellate growth. After testing multiple different antibiotics, we grew *Hartaetosiga kampana* cells for several passages in 1:100 RS medium with 50 μg/ml streptomycin (S9137, Sigma-Aldrich) until no further change in bacterial density or diversity was observed. Meanwhile, *Oinochoeca dipylis* cells were initially treated for several times with a sterilized antibiotic solution of 50 μg/ml streptomycin diluted in 1:100 RS medium and subsequently diluted into fresh medium (1:100 RS medium) with a final concentration of 50 μg/ml kanamycin (K1377-5G, Sigma-Aldrich) repeatedly until the limitation of bacterial density and diversity was the optimal.

*Salpingoeca kylix* (BEAP0192, C3C) was isolated in April 2014 from a salt evaporation pond in Eilat, Israel (GPS: 29.629264, 35.001874). The sample was enriched in a hypersaline medium (150 ppt) containing NaCl, KCl, MgCl_₂_.6H₂O, MgSO₄.7H₂O, and CaCl_₂_.2H₂O (<u>Park</u> <u>2012</u>). Several rounds of dilution were performed into the same medium to remove contaminating eukaryotes. The strain is maintained at 23 °C under a 12:12 light:dark cycle in the same medium, supplemented with a single grain of rice.

*Daedalina diplosolina* (BEAP0272, Gre159) was collected in February 2023 from a mangrove habitat in Grenada (GPS: 12.2265547, −61.6102777). For enrichment, 2 ml of sediment was added to 3 ml of freshwater RS medium (1:50 P5:distilled water) in a 6-well plate. After 4 days, the culture was transferred to 20 ml of fresh medium and serially diluted to eliminate non-target eukaryotes. The strain was initially maintained at 23 °C under a 12:12 light:dark cycle. Subsequently, cultures are maintained in RS medium (1:50 P5:tap water) at 18°C with natural light, which consistently supports improved growth.

*Hippolytella eleftheri* (BEAP0279, FR2326) was isolated in May 2023 from the bank of the Fraser River in British Columbia, Canada (GPS: 49.1101852, −123.1321428), from sediment and algae. A single cell was isolated into 200 μL of Pratt’s freshwater medium (‘Pratt’s Freshwater Medium’ 2009) supplemented with 0.025% wheat grass extract. The culture is now maintained in freshwater RS medium (1:500 P5:distilled water) at 17 °C in darkness. We limited the diversity and density of bacterial prey in this culture by treating several passages in 1:500 RS medium with 40 μg/ml Ampicillin (6800, Condalab) until no further change in bacterial density or diversity was observed.

*Stagondoeca pyriformis* (Carr et al., 2017) (BEAP0306, CHO_A3) was re-isolated during this study at July 2023 from coastal sediment at Vigna sul Mar beach, Ferrara, Italy (GPS: 44.719278, 12.243972). Sampling was conducted at approximately 4 m depth. Cultures were initiated in marine Cerophyl medium (MWC: 5 ml Cerophyl solution in 45 ml filtered marine water) and cleaned from contaminating eukaryotes via serial dilutions. The strain is now maintained at 17 °C under a natural light cycle using RS medium (1:100 P5:P4).

*Salpingoeca rosetta* strain Px1 (ATCC PRA-366) was obtained from the laboratory of Iñaki Ruiz-Trillo and Elena Casacuberta (Institute of Evolutionary Biology, Spain). The culture is maintained in RS medium (P5, 1:100) at 17 °C in the dark.

*Salpingoeca kvevrii* (ATCC 50929) and *Salpingoeca urceolata* (ATCC 50560) were obtained as frozen stocks from the lab of Nicole King (University of California, Berkeley). Both strains were revived in RS medium (1:100 P5:P4) and are maintained with their co-isolated mixed bacterial prey at 17 °C in darkness.

*Salpingoeca tuba* (BEAP0445) (Kent, 1882) was re-isolated during this study in April 2025 from a marine splash pool at Pointe des Chats, Groix Island, France. The sample was enriched by replacing the supernatant with fresh artificial seawater (ASW) supplemented with 0.1% SWC medium. The strain is maintained in marine RS medium (1:100 P5:P4) at 18 °C under a natural light cycle as a mixed culture containing multiple bacterial and eukaryotic species.

### Morphological analysis

**<u>Differential interference contrast (DIC) microscopy:</u>** Cells from all strains were seeded directly onto the glass surface of imaging dishes (150680 or 80807, Thermo Fisher Scientific). Imaging, including time-lapse video acquisition, was performed using an Axio Observer 7 inverted microscope equipped with a Plan-Apochromat 63x/1.4NA DIC M27 oil-immersion objective, Hamamatsu HDCamC13440-20CU camera and ZEN blue image acquisition software. Images and time-lapses were processed using Fiji software (ImageJ) (Schneider et al., 2012). For the scraping experiments presented in Supplementary Figure 4B, *Daedalina diplosolina* cells were seeded onto glass-bottom imaging dishes (150680, Thermo Fisher Scientific) and allowed to grow for 1 week before imaging. DIC images were acquired immediately before and immediately after mechanical scraping to quantify the proportions of thecate cells, naked cells and detached cells containing only the upper part of the theca. In the first experiment (N1), 153 cells were counted before scraping and 152 cells after scraping. Following optimization of the culture conditions to RS medium (1:50 P5:tap water) at 18°C, which supported improved growth, the experiment was repeated (N2) using the same procedure, with 241 cells counted before scraping and 242 cells after scraping.

**<u>Confocal microscopy</u>:** To visualize the cytoskeletal structures including the collar of microvilli, filopodia and flagella, cells were stained with a combination of phalloidin (F-actin), β-Tubulin antibody (flagella/microtubules) and Hoechst dye (DNA). Cells were seeded on coverslips either pre-treated with poly-L-lysine or left untreated, depending on the adherence efficiency of each strain, washed once with washing buffer, and fixed with a solution of 0.25% glutaraldehyde and 1% formaldehyde in washing buffer for 5 minutes. After a 5-minute wash with the same buffer, cells were blocked and permeabilized in a solution containing 1% BSA and 0.3% Triton X-100 in washing buffer for 1 hour. Cells were then incubated overnight at 4 °C with 1:50 β-tubulin mouse antibody (E7-S, Hybridoma Bank) diluted in the permeabilization/blocking buffer. The following day, cells were washed twice with washing buffer and incubated for 1 hour at room temperature with 1:200 Alexa Fluor 568-conjugated anti-mouse antibody (A11061, Molecular Probes), followed by two additional washes. Next, cells were incubated for 30 minutes at room temperature with 1:50 phalloidin 488 (A12379, Thermo Fisher Scientific) in washing buffer. DNA was stained with 1:2000 Hoechst 33258 (H3570, Life Technologies) in washing buffer. Coverslips were mounted in ProLong Gold antifade reagent (P36934, Thermo Fisher Scientific). The composition of the washing buffer varied depending on the strain: artificial seawater was used for marine species (*Hartaetosiga kampana*), while PBS was used for freshwater strains (*Hippolytella eleftheri* and *Daedalina diplosolina*).

To visualize membranes (DiOC6(3)) and extracellular structures (α-poly-L-lysine conjugated to Alexa Fluor 546 dye), we followed a protocol adapted from Colin et al. (2017) (Colin et al., 2017). PLL546 was prepared by conjugating poly-L-lysine (Sigma-Aldrich, cat. no. P5899) to Alexa Fluor 546 succinimidyl ester (Invitrogen, cat. no. A20002), as suggested by Colin et al. (2017). Cells were seeded on coverslips pre-coated with poly-L-lysine, washed once with washing buffer, and fixed for 5 minutes with a solution of 0.25% glutaraldehyde and 1% formaldehyde in washing buffer. After an additional wash, cells were incubated with 0.1 mg/ml PLL546, diluted in washing buffer, for 15 minutes at room temperature. This was followed by a 5-minute wash and incubation with 1.5 μM DiOC6(3) (D273, Invitrogen) for 30 minutes at room temperature, also diluted in washing buffer. DNA was then stained with 1:2000 Hoechst 33258 in washing buffer, and coverslips were mounted using ProLong Gold antifade reagent. The washing buffer composition varied depending on the strain. Artificial seawater was used for marine species like *Hartaetosiga kampana*, *Oinochoeca dipylis*, *Salpingoeca rosetta*, *Salpingoeca kvevrii*, *Salpingoeca urceolata* and *Stagondoeca pyriformis*. PBS was used for freshwater strains like *Daedalina diplosolina* and *Hippolytella eleftheri*. Finally, a hypersaline medium was used as washing buffer for *Salpingoeca kylix*.

The observation of the samples was carried out at the Advanced Light Microscopy Unit (ALMU) located at the Centre for Genomic Regulation (CRG) in Barcelona, using a Zeiss LSM 980 confocal microscope equipped with Airyscan 2 super-resolution (405, 488, 561, 639 nm lasers) and a 63x lens. Digital images were later processed using Fiji software (Schneider et al., 2012).

We used poly-L-lysine staining as a general strategy to label extracellular material, given that the molecular composition of extracellular structures remains unknown in most of the species examined. Poly-L-lysine, due to its positive charge, adheres non-specifically to negatively charged surfaces, allowing broad detection of surface-associated or secreted extracellular features. For the extracellular morphological analysis, each species was imaged in three independent replicates, with ten representative images of thecate or empty-theca structures acquired per replicate. The general extracellular morphology was found to be consistent across all replicates and observed cells.

**<u>3D-reconstruction analysis</u>:** We obtained serial super-resolution sections of thecate cells, as well as empty thecae stained with DiOC6(3), Hoechst and α-poly-L-lysine conjugated to Alexa Fluor 546 dyes using a Zeiss LSM 980 microscope equipped with Airyscan 2. The 3D reconstruction analysis was carried out at the Advanced Light Microscopy Unit (ALMU) located at the Centre for Genomic Regulation (CRG) in Barcelona, using Imaris Cell Imaging Software. The 3D-reconstruction analysis incorporated all staining channels (Hoechst, DiOC6(3), and PLL546). Additional 3D-surface reconstruction was conducted using solely the PLL546 channel.

<u>TEM observation after High-Pressure Freezing and Freeze Substitution for the strains</u> *<u>Hartaetosiga kampana</u>* <u>and *Oinochoeca dipylis*</u>: Transmission electron microscopy sample preparation was performed in the Electron Cryomicroscopy Unit at the Scientific and Technical Centers of the University of Barcelona. Briefly, samples were concentrated by centrifugation (2000 x g, 5 minutes) and resuspended in 20% BSA in artificial sea water. Most of the supernatant was removed and the concentrated cells cryo-immobilized using a Leica HPM100 high-pressure freezer (Leica Microsystems, Vienna, Austria). Samples were freeze-substituted in pure acetone containing 2% (w/v) osmium tetroxide and 0.1% (w/v) uranyl acetate at −90 °C for 72 hours in an EM AFS2 (Leica Microsystems, Vienna, Austria). Later, they were warmed up to 4 °C at a 5 °C/h slope, kept at 4 °C for 2 hours, and transferred to room temperature and kept for 2 hours in darkness. Samples were washed in acetone at room temperature, infiltrated in increasing concentrations of Epon-812 resin in acetone until purely Epon-812. Then, they were embedded and polymerized in Epon-812 at 60 °C for 48 hours. Ultrathin sections of 60 nm were obtained with a UC6 ultramicrotome (Leica Microsystems, Vienna, Austria) and placed on Formvar-coated copper grids. Sample sections were stained with 2% (w/v) uranyl acetate for 30 minutes, lead citrate for 5 minutes and examined in a TEM Jeol JEM 1010 (Gatan, Japan) equipped with a tungsten cathode. Images were acquired at 80 kV with a 1k x 1k CCD Megaview III camera.

**<u>SEM observation after critical point drying for *Hartaetosiga kampana*</u>**: Scanning Electron Microscopy preparation for the strains was performed on the TEM-SEM Electron Microscopy Unit from Scientific and Technological Centers (CCiTUB), University of Barcelona. Sample was deposited on poly-L-lysine glass coverslips and fixed in a solution consisting of 2.5% glutaraldehyde in 0.1 M phosphate buffer (pH 7.4), post-fixed in osmium tetroxide (1%) in the same phosphate buffer, dehydrated in graded alcohol and processed for critical point drying using an Emitech K850. The sample was covered with a thin carbon film in order to improve their electrical conductivity. The sample was observed with a Jeol JSM-7001F (Jeol, Japan) operated at 10 kV.

**<u>SEM observations after critical point drying for *Daedalina diplosolina, Salpingoeca tuba*</u> <u>and *Hippolytella eleftheri*</u>**: Cells were fixed with 2.5% glutaraldehyde for 3 hours at room temperature and then seeded on membranes with pores sized 0.8 μm (WHA10417301, MERCK Chemicals and Life Science) via filtering. The samples were dehydrated in a graded ethanol series and critical-point dried with liquid carbon dioxide in a Leica EM CPD300 unit (Leica Microsystems, Austria). The dried filters were mounted on stubs with colloidal silver and then were sputter-coated with gold (*D. diplosolina* and *H. eleftheri*) or iridium (*S. tuba*) in a Q150R S (Quorum Technologies, Ltd.) and observed either with a Hitachi S3500N scanning electron microscope (Hitachi High Technologies Co., Ltd, Japan) at an accelerating voltage of 5 kV or with a Hitachi SU8600 field emission scanning electron microscope (Hitachi High Technologies Co., Ltd., Japan) in the Electron Microscopy Service of the Institute of Marine Science (ICM-CSIC), Barcelona.

### 18S rRNA gene sequencing

*Stagondoeca pyriformis* (BEAP0306, CHO_A3) cells were harvested by centrifugation at 15,000 rpm for 10 minutes. Genomic DNA was extracted using the MasterPure kit (MC85200, Epicentre), following the manufacturer’s standard protocol. The 18S rRNA gene was amplified using the universal eukaryotic primers EukA (5′-AACCTGGTTGATCCTGCCAGT-3′) and EukB (5′-TGATCCTTCTGCAGGTTCACC-3′) (Countway et al., 2005) with the KAPA HiFi HotStart ReadyMix PCR Kit (KK2601, Roche). Amplification products were visualized on a 1% agarose gel. Bands of the expected size were purified by gel extraction using NZY gel pure kit (MB01103, NZYTECH LDA). The purified amplicons were Sanger sequenced at Macrogen (Seoul, Korea) using the four primers, EukA, EukB, 528F-Carr (5’-CGGTAATTCCAGCTCC-3’) and 1055R-Carr (5’-CGGCCATGCACCACC-3’). The reads obtained from sequencing were assembled using Tracy (Rausch et al., 2020). The consensus sequence was compared via BLAST to the PR2 database, where it showed a 99.13% identity to *Stagondoeca pyriformis* (KT757499) (Carr et al., 2017). After further evaluation, the minor differences were attributed to strain-level variation and potential PCR or sequencing errors, thereby confirming species identity.

*Salpingoeca tuba* cells were scraped and harvested by centrifugation at 3300g for 10 minutes. Supernatant was removed and cells were incubated with DNAzol DIRECT (MRC, Cat No: DN131) for 15 min with agitation. The 18S rRNA gene was amplified using choano-specific primers Cs3F (5’AAAGATTAAGCCATGCATGTCTAAG 3’) and Cs1R (5’CAGGGACGTAATCAACGCGA 3’) with Q5 DNA polymerase (NEB). Annealing was performed at 63°C and extension was done for 1 min, 40 cycles were used. The PCR product was purified on a gel and sequenced with the same primers. The sequence obtained was matching the one previously available for that species (HQ026774) .

The 18S/28S rRNA sequences for the remaining strains were extracted from the contigs assembled with either Trinity or rnaSPAdes during the transcriptomic analysis.

### Transcriptomic analysis

**<u>RNA extraction:</u>** To extract and sequence total RNA from *Hartaetosiga kampana*, *Oinochoeca dipylis*, *Salpingoeca kylix*, *Daedalina diplosolina* and *Hippolytella eleftheri* strains, we employed a similar experimental approach as Richter and colleagues (Richter et al., 2018). Prior to RNA isolation, all strains were grown in large batches using 75 cm^2^ Rectangular Canted Neck Cell Culture Flasks with Vented Caps (353136, Corning Life Sciences), each containing 50 ml of medium (2 for each strain). Total RNA from all strains was isolated by using the RNAqueous kit (AM1912, ThermoFisher Scientific), following the manufacturer’s protocol with modifications described by (Richter et al., 2018) to improve RNA yield and avoid degradation. RNA concentration was measured using a NanoDrop One/One C UV-Vis Microvolume Spectrophotometer (ThermoFisher Scientific). We digested genomic DNA using the TURBO DNA-free kit (AM1907, ThermoFisher Scientific) according to the manufacturer’s instructions and removed DNase using DNase Inactivation Reagent. The quality evaluation of extracted total RNA was carried out at the CRG Genomics Core Facility in Barcelona. In detail, the quality of extracted total RNA was assessed by using Bioanalyzer 2100 RNA Pico chips (*Hartaetosiga kampana*, *Oinochoeca dipylis* and *Daedalina diplosolina*) or Nano chips (*Salpingoeca kylix* and *Hippolytella eleftheri*) (Agilent Technologies). The library preparation and sequencing was performed either at the CRG Genomics Core Facility in Barcelona, Spain (for strains *Hartaetosiga kampana*, *Oinochoeca* dipylis, *Salpingoeca kylix* and *Hippolytella eleftheri*) or Macrogen in Seoul, Korea (*Daedalina diplosolina*). To prepare stranded libraries, the NEBNext Ultra II Directional RNA Library Prep Kit Total RNA was used, applying double the normal rounds of PolyA selection prior to cDNA synthesis to reduce the excess of bacterial RNA. Finally, the RNA was sequenced by using Illumina NextSeq 2000 (2×150 bp).

***<u>De novo</u>* <u>transcriptome assembly</u>:** The quality of the RNA sequencing data was evaluated using FastQC v0.11.9 (https://www.bioinformatics.babraham.ac.uk/projects/fastqc) and sequence purity was assessed with phyloFlash v3.4 (Gruber-Vodicka et al., 2020), which estimates the taxonomic composition of metagenomic datasets based on 16S/18S rRNA genes. Removal of sequencing artifacts and adapter contamination was performed using fastp v0.23.2 (Chen et al., 2018) with the parameters: --low_complexity_filter, --cut_front, --cut_tail, --cut_right, --cut_front_window_size 1, --cut_tail_window_size 1, --cut_right_window_size 4, --cut_mean_quality 5, --trim_front1 12, --trim_front2 12, --trim_poly_g, --trim_poly_x, --adapter_sequence=AGATCGGAAGAGCACACGTCTGAACTCCAGTCA, and --adapter_sequence_r2=AGATCGGAAGAGCGTCGTGTAGGGAAAGAGTGT. Transcriptomes were assembled *de novo* using Trinity v2.14.0 (Grabherr et al., 2011), applying the strand-specific parameter --SS_lib_type RF.

Following assembly, potential contaminants were identified through phyloFlash analysis and BLASTn searches of Trinity-assembled contigs. Briefly, 16S/18S rRNA sequences recovered by phyloFlash were used as queries against the corresponding Trinity assemblies and matching contigs were subsequently compared against EukRibo or NCBI databases to assess their taxonomic identities. Based on these results, decontamination was performed using BLASTn against a curated set of potential contaminant genomes. The distributions of BLASTn percentage identity and alignment length were examined to guide the selection of strain-specific thresholds. For *Hartaetosiga kampana*, sequences showing ≥98% identity and alignment length >100 bp to *Cutibacterium acnes* (ASM673938v1) were excluded. For *Salpingoeca kylix*, contamination was restricted to insect sequences. Because no genome was available for the exact contaminant species, comparisons were made against several closely related insect genomes (ASM2641340v1, ASM2641346v1, GCA_029228625.1, GCA_965233325.1, GCA_965233655.1, GCF_014529535.1), and contigs with ≥97% identity over >100 bp were removed. For the remaining species, *Oinochoeca dipylis*, *Daedalina diplosolina* and *Hippolytella eleftheri*, although bacterial 16S rRNA sequences were detected by phyloFlash, no contaminant contigs were identified through BLASTn searches against the corresponding bacterial reference genomes.

To minimize cross-sample contamination within each sequencing run, WinstonCleaner (https://github.com/kolecko007/WinstonCleaner) was employed using default parameters on sets of samples that were sequenced together. However, the sample *Hippolytella eleftheri* was heavily cross-contaminated by another culture, a mixed sample containing a choanoflagellate and an amoeba (Cutosea species), that had been sequenced in the same run. To further improve decontamination of *Hippolytella eleftheri*, we applied an additional BLASTn filtering step. Contigs showing sequence identity ≥94% to the contaminant dataset, with alignment lengths exceeding 100 bp, were removed. In addition, because the raw reads intended for public deposition still contained contaminant reads derived from the second choanoflagellate and the amoeba, both of which are the subject of ongoing independent analysis, the reads were further filtered against the cleaned curated transcriptome. More specifically, reads were mapped using Bowtie 2 v2.4.4 (Langmead and Salzberg, 2012) in end-to-end and very-sensitive mode, and only properly paired reads with high-confidence unique alignments (MAPQ ≥ 40) were retained using SAMtools v1.13 (Li et al., 2009). The filtered read set was subsequently remapped to the curated transcriptome to validate the filtering procedure.

Protein sequences were predicted using TransDecoder v5.5.0 (Haas, 2017; Haas et al., 2013), applying the -S option to maintain strand specificity. Functional domain annotation was conducted with InterProScan v5.56-89.0 (Jones et al., 2014), using the options --disable-precalc and --applications Pfam, Phobius. Transcriptome completeness was assessed by identifying conserved single-copy orthologs through BUSCO v5.3.2, using the Eukaryota odb10 dataset (Manni et al., 2021; Simão et al., 2015). For all strains, transcriptomes were also assembled independently with rnaSPAdes v4.1.0 (Bankevich et al., 2012) (using strand-specific parameters, --ss rf), after which the same decontamination, annotation and completeness assessment pipeline was applied. This approach allowed cross-validation of assemblies and retrieval of ribosomal RNA genes for use in subsequent phylogenetic analyses.

### Ribosomal gene phylogenies and phylogenomic analysis

**<u>PhyloFisher:</u>** The predicted proteomes of *Hartaetosiga kampana*, *Oinochoeca dipylis*, *Salpingoeca kylix*, *Daedalina diplosolina* and *Hippolytella eleftheri* were processed following the standard PhyloFisher (Tice et al., 2021)) pipeline for phylogenomic assembly. Briefly, the pipeline includes homolog identification, single-gene tree reconstruction, ortholog selection, multiple sequence alignment, trimming and concatenation into a supermatrix. The resulting alignment consisted of 53 taxa: the 5 newly assembled transcriptomes, 26 additional choanoflagellate species retrieved from the EukProt database (Richter et al., 2022) and 22 outgroup holozoans (9 protists and 13 Metazoa) already included in the PhyloFisher dataset. Maximum likelihood phylogenetic analysis was performed using IQ-TREE v3.0.0 (Wong et al., 2026). The best fitting model (LG+F+I+R7) was automatically selected based on the Bayesian Information Criterion (BIC). Statistical support was assessed using 1000 ultrafast bootstrap replicates and trees were visualized using iToL (Letunic and Bork, 2021).

**<u>Tree based on rRNA gene sequences:</u>** A concatenated rRNA gene dataset consisting of 4260 aligned nucleotide sites (1525 from 18S, 135 from 5.8S and 2600 from 28S) was assembled for 52 choanoflagellate taxa and 10 outgroups (2 protists and 8 Metazoa). Sequences were retrieved from GenBank or extracted from transcriptome, genome and single-amplified-genome assemblies (accessions and contigs in **Supplementary Table 1**). Sequences were aligned manually according to a secondary structure model (Ben Ali et al., 2001) and manually trimmed. Maximum likelihood analysis was performed using RAxML-HPC (Stamatakis, 2014) under the GTRGAMMA model. Branch support was evaluated using the rapid bootstrap algorithm with 1000 replicates and the resulting tree was visualized in iToL (Letunic and Bork, 2021).

### Taxonomic summary

Eukaryota, Amorphea, Obazoa, Opisthokonta, Holozoa, Choanozoa, Choanoflagellatea, Craspedida

The following taxa are assigned to the Order Craspedida Cavalier-Smith 1997:

Hartaetosiga kampana sp. nov.

The species *Hartaetosiga kampana* is a member of the genus *Hartaetosiga* Carr, Richter and Nitsche 2017.

**Diagnosis:** Cells possess an ovoid cell body approximately 3.6 µm in width and 4.5 µm in length (n = 20). The collar is around 5.3 µm in length and surrounds a single anterior flagellum measuring 6.8 ± 2.2 µm in length (n = 20). The cell body is surrounded by a close fitting glycocalyx coat that partially covers the collar of microvilli and can extend into a peduncle.

**Etymology:** The species name is derived from the Greek word kampana which refers to the bell-like morphology of motile cells we observed during the scanning electron microscopy experiments.

**Type strain:** *Hartaetosiga kampana* (BEAP0001, CRO2OCHO1, RCC7815, CCAP (deposited))

**Type locality:** Shallow marine lagoon Malo Jezero on Mljet Island, Croatia.

**Type material**: The name-bearing type (an hapantotype) is a culture containing *Hartaetosiga kampana* deposited in the Roscoff Culture Collection with accession number RCC7815 and in the Culture Collection of Algae and Protozoa with accession number (deposited). An SEM stub of fixed cells from this culture is deposited at the Marine Biological Reference Collections (CBMR) at the Institut de Ciències del Mar (ICM-CSIC, Barcelona, Spain) under the catalog/accession number (submitted) (Guerrero et al., 2016). This material also contains uncharacterized prokaryotic species, which do not form part of the hapantotype.

**Isolated by**: Luis Javier Galindo

**Gene sequence:** The rRNA gene sequence from *Hartaetosiga kampana* was deposited in GenBank with accession number QB248917.

**ZooBank registration**: Described under the Zoological Code; ZooBank registration will be performed following peer review.

***Stagondoeca* emend.**

**Diagnosis:** Small, uninucleate protists with a single, centrally positioned, anterior flagellum, which is surrounded by a collar of long, actin-supported microvilli. Phagotrophic. Cell body enclosed in a robust organic three-lobed shaped theca from which a pedicel extrudes.

**Type species**: Stagondoeca pyriformis

Oinochoeca gen. nov.

**Diagnosis:** Cells measure approximately 5 µm in length and possess a single anterior flagellum surrounded by a collar of actin-based microvilli. The organisms are phagotrophic. During part of the life cycle, cells are enclosed within an organic three-lobed shaped theca, from which a pedicel extends. Thecate cells are additionally characterized by the presence of one or more organic sheath-like layers surrounding the cell body. Additional non-thecate swimming and crawling morphotypes are also observed in the culture.

**Etymology:** The genus name is derived from the Greek word oinochoe that refers to a wine jug and describes the shape of the theca. Oinochoeca is feminine.

**Type species:** Oinochoeca dipylis

**ZooBank registration**: Described under the Zoological Code; ZooBank registration will be performed following peer review.

Oinochoeca dipylis sp. nov.

**Diagnosis:** Cell body is ovoid in shape, measuring around 5.3 µm in length and 3.6 µm in width (n = 21). A collar, approximately 6.3 µm (n = 21) in length, surrounds a flagellum of 5.5 ± 0.8 µm in length (n = 16). Attached cells generate a three-lobed theca, which extends into a pedicel, and are characterized by an additional sheath that partially covers the collar of microvilli in a close-fitting way. Additional crawling cells lacking a visible theca are also observed in culture, measuring around 3.4 µm in width and 4.1 µm in length (n = 5), possessing a short collar of microvilli measuring approximately 1.3 µm in length, an anterior flagellum measuring 7.0 ± 1.8 µm in length and posterior filopodia.

**Etymology**: The species name is derived from the Greek word dipylis (“double-gated”), referring to the presence of an additional sheath.

**Type strain:** *Oinochoeca dipylis* (BEAP0002, LJO2, RCC7816, CCAP (deposited))

**Type locality:** Playa del Janubio in Lanzarote, Canary Islands, Spain.

**Type material**: The name-bearing type (an hapantotype) is a culture containing *Oinochoeca dipylis* and mixed bacterial species deposited in the Roscoff Culture Collection with accession number RCC7816 and in the Culture Collection of Algae and Protozoa with accession number (deposited). A resin-embedded sample block and TEM grids containing ultrathin sections of cells are deposited at the Marine Biological Reference Collections (CBMR) at the Institut de Ciències del Mar (ICM-CSIC, Barcelona, Spain) under the catalog/accession number (submitted) (Guerrero et al., 2016). This material also contains uncharacterized prokaryotic species, which do not form part of the hapantotype.

**Gene sequence:** The rRNA gene sequence from *Oinochoeca dipylis* was deposited in GenBank with accession number QB248919.

**Isolated by:** Luis Javier Galindo

**ZooBank registration**: Described under the Zoological Code; ZooBank registration will be performed following peer review.

Salpingoeca kylix sp. nov.

The species *Salpingoeca kylix* is a member of the genus *Salpingoeca* James-Clark 1866.

**Diagnosis:** The roughly spherical-shaped cell body is approximately 3.9 µm in width and 4.8 µm in length (n = 20). The collar is around 5.7 µm in length and surrounds a flagellum measuring 21.7 ± 9.3 µm in length (n = 20). Attached cells produce a cup-shaped theca which covers the bottom edge of the cell body and extends into a long pedicel. The cell body is enclosed in an additional sheath that possesses a neck.

**Etymology:** The species name is derived from the Greek word kylix, referring to the morphology of the extracellular theca. Kylix denotes a shallow drinking cup and describes the cup-like shape of the theca.

**Type strain:** *Salpingoeca kylix* (BEAP0192, C3C, CCAP (deposited))

**Type locality:** Hypersaline salt evaporation pond in Eilat, Israel.

**Type material**: The name-bearing type (an hapantotype) is a culture containing *Salpingoeca kylix* and mixed bacterial species deposited in the Culture Collection of Algae and Protozoa with accession number (deposited).

**Isolated by**: Yana Eglit

**Gene sequence:** The rRNA gene sequence from *Salpingoeca kylix* was deposited in GenBank with accession number QB248918.

**ZooBank registration**: Described under the Zoological Code; ZooBank registration will be performed following peer review.

Daedalina gen. nov.

**Diagnosis:** Cells measure approximately 7 µm in length and possess a single anterior flagellum surrounded by a collar of actin-based microvilli. The organisms are phagotrophic. During part of the life cycle, cells are enclosed within an organic tubular-shaped two-parted theca, from which a pedicel may extend. Cell division proceeds by an inverted division mode, with daughter cells separating flagellar pole to flagellar pole. Additional swimming and crawling morphotypes are observed in culture. Crawling cells lack a visible theca and are characterized by a long anterior flagellum, a reduced collar of microvilli, and prominent posterior filopodia.

**Etymology:** The genus name is derived from Daedalus, the craftsman of Greek mythology, referring to the intricate architecture of the theca. Daedalina is feminine.

**Type species:** Daedalina diplosolina

**ZooBank registration**: Described under the Zoological Code; ZooBank registration will be performed following peer review.

Daedalina diplosolina sp. nov.

**Diagnosis:** Cell body elongate, measuring around 7.4 µm in length and 3.6 µm in width (n = 21). The collar of actin-based microvilli measures approximately 6.1 µm in length (n = 19), and the anterior flagellum measures 7.4 ± 2.7 µm in length (n = 20). Crawling cells measure around 5.8 µm in length and 3.4 µm in width (n = 4), possess a reduced collar measuring approximately 0.9 µm in length, an anterior flagellum measuring 25.9 ± 0.8 µm in length, and produce posterior filopodia.

**Etymology:** The species name is derived from the Greek words diploos (“double”) and solen (“tube”), referring to the two-parted tubular theca.

**Type strain**: *Daedalina diplosolina* (BEAP0272, Gre159, CCAP (deposited))

**Type locality:** Freshwater mangrove habitat, Grenada, Caribbean.

**Type material**: The name-bearing type (an hapantotype) is a culture containing *Daedalina diplosolina* and mixed bacterial species deposited in the Culture Collection of Algae and Protozoa with accession number (deposited). An SEM stub of fixed cells from this culture is deposited at the Marine Biological Reference Collections (CBMR) at the Institut de Ciències del Mar (ICM-CSIC, Barcelona, Spain) under the catalog/accession number (submitted) (Guerrero et al., 2016). This material also contains uncharacterized eukaryotic and prokaryotic species, which do not form part of the hapantotype.

**Isolated by**: Daryna Zavadska

**Gene sequence:** The rRNA gene sequence of *Daedalina diplosolina* was deposited in GenBank under accession number QB248920.

**ZooBank registration**: Described under the Zoological Code; ZooBank registration will be performed following peer review.

Eukaryota, Amorphea, Obazoa, Opisthokonta, Holozoa, Choanozoa, Choanoflagellatea

The following taxon is not assigned to any currently described order within the class Choanoflagellatea Cavalier-Smith 1998:

Hippolytella gen. nov.

**Diagnosis:** Cells measure approximately 5 µm in length and possess a single anterior flagellum surrounded by a collar of actin-based microvilli and are phagotrophic. Cells appear naked, lacking a silicified lorica or other rigid extracellular covering and possess thin actin-rich filopodia extending across the cell body. Both swimming and substrate-associated crawling behaviours are observed in culture.

**Etymology:** The name is derived from Hippolyta, the legendary queen of the Amazons in Greek mythology, with the diminutive suffix -ella. The name refers to the characteristic depiction of Amazons wearing scythian trousers in classical Greek art and alludes to the absence of the lorica, the characteristic external “dress” of its closest relatives. *Hippolytella* is feminine.

**Type species**: Hippolytella eleftheri

**ZooBank registration**: Described under the Zoological Code; ZooBank registration will be performed following peer review.

Hippolytella eleftheri sp. nov.

**Diagnosis:** Cells are ovoid in shape, measuring approximately 4.9 µm in length and 4.4 µm in width (n = 20). A collar of microvilli measuring around 3.3 µm in length surrounds a single anterior flagellum measuring 7.1 ± 2.7 µm in length (n = 20). Cells appear naked, lacking a silicified lorica or other rigid extracellular covering. Thin actin-rich filopodia extend across the cell body and may reach up to four times the cell length. Cells display both swimming and substrate-associated crawling behaviours.

**Etymology**: The species name is derived from the Greek *eleutheria*, meaning “freedom” or “liberty”, referring to the absence of a silicified lorica.

**Type strain**: *Hippolytella eleftheri* (BEAP0279, FR2326, CCAP (deposited))

**Type locality:** Freshwater, bank of the Fraser River, British Columbia, Canada.

**Type material**: The name-bearing type (an hapantotype) is a culture containing *Hippolytella eleftheri* and mixed bacterial species deposited in the Culture Collection of Algae and Protozoa with accession number (deposited). An SEM stub of fixed cells from this culture is deposited at the Marine Biological Reference Collections (CBMR) at the Institut de Ciències del Mar (ICM-CSIC, Barcelona, Spain) under the catalog/accession number (submitted) (Guerrero et al., 2016). This material also contains uncharacterized prokaryotic species, which do not form part of the hapantotype.

**Isolated by:** Vasily Zlatogursky

**Gene sequence:** The rRNA gene sequence from *Hippolytella eleftheri* was deposited in GenBank with accession number QB248921.

**ZooBank registration**: Described under the Zoological Code; ZooBank registration will be performed following peer review.

This publication (work) will be registered with ZooBank following peer review.

## Acknowledgements

Regarding *Salpingoeca kylix* we thank Aharon Oren (the Hebrew University of Jerusalem) for collecting the samples, and Salt of the Earth Ltd. for allowing access to the Eilat salterns. Moreover, we would like to thank Maria Carlos Oliveira for expert assistance with figure preparation, Àlex Gàlvez-Morante for providing the images of *Acanthocorbis camarensis* and Meritxell Antó for assisting in the molecular identification of *Stagondoeca pyriformis* (BEAP0306, CHO_A3). We would also like to thank Thibaut Brunet for providing the *Salpingoeca tuba* culture along with valuable insights and feedback regarding this work.

## Data Availability

Supplementary material (including videos), transcriptome assemblies, protein predictions, sequence alignments, phylogenetic trees, and additional data are available on FigShare at the DOI: 10.6084/m9.figshare.34737594. Raw transcriptome reads are available via NCBI BioProject PRJNA1085734. Ribosomal gene sequences are available in GenBank with the identifiers QB248917-QB248921 and QB212100-QB212101. SEM stubs of fixed cells are deposited at the Marine Biological Reference Collections (CBMR) at the Institut de Ciències del Mar (ICM-CSIC, Barcelona, Spain) under the catalog/accession numbers (submitted) (Guerrero et al., 2016).

---

**Supplementary Figure 1:**
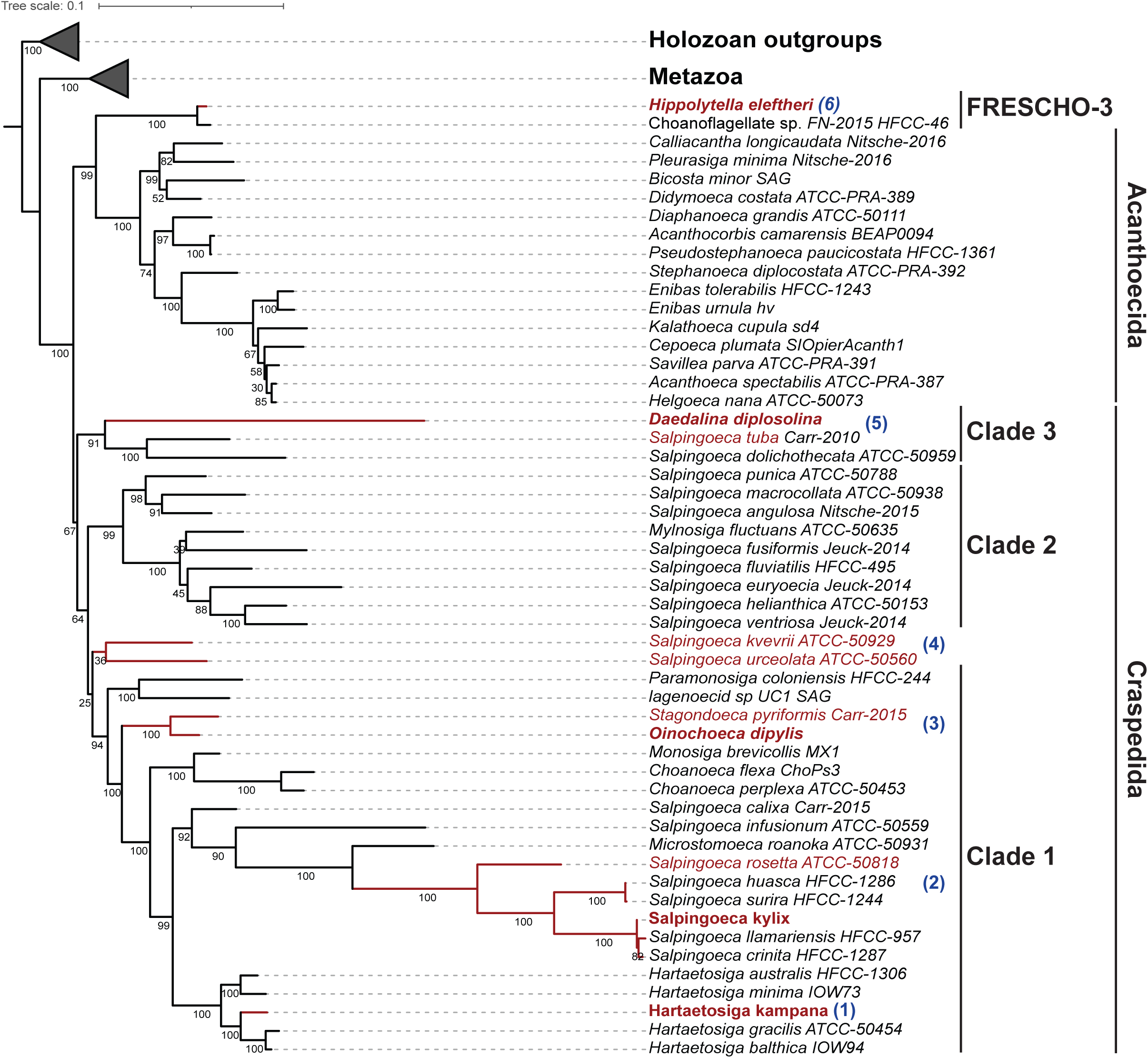
Phylogenetic reconstruction based on concatenated rRNA gene sequences (18S, 5.8S and 28S). The dataset included 52 choanoflagellate taxa and 10 outgroup holozoans (2 protists and 8 Metazoa). Maximum likelihood inference was performed with RAxML (Stamatakis, 2014). Bootstrap values are shown below the internal branches. *Hartaetosiga kampana* clusters within the genus *Hartaetosiga* with strong support, while *Salpingoeca kylix* branches within a strongly-supported group of *Salpingoeca* species that includes *S. rosetta*. *Oinochoeca dipylis* is sister to *Stagondoeca pyriformis*. *Daedalina diplosolina* is part of Clade 3, branching sister to the clade containing *Salpingoeca dolichothecata* and *S. tuba*. The close relationship between *S. kvevrii* and *S. urceolata* is also recovered, although with low support. *Hippolytella eleftheri* is sister to the choanoflagellate FN-2015, together forming a well-supported sister group to the Acanthoecida. Outgroups are collapsed for clarity. The species whose extracellular structures were stained and compared in this study are highlighted in red, the species isolated as part of this study are in bold, and blue labels indicate the order of morphological comparisons in the text.

**Supplementary Figure 2:**
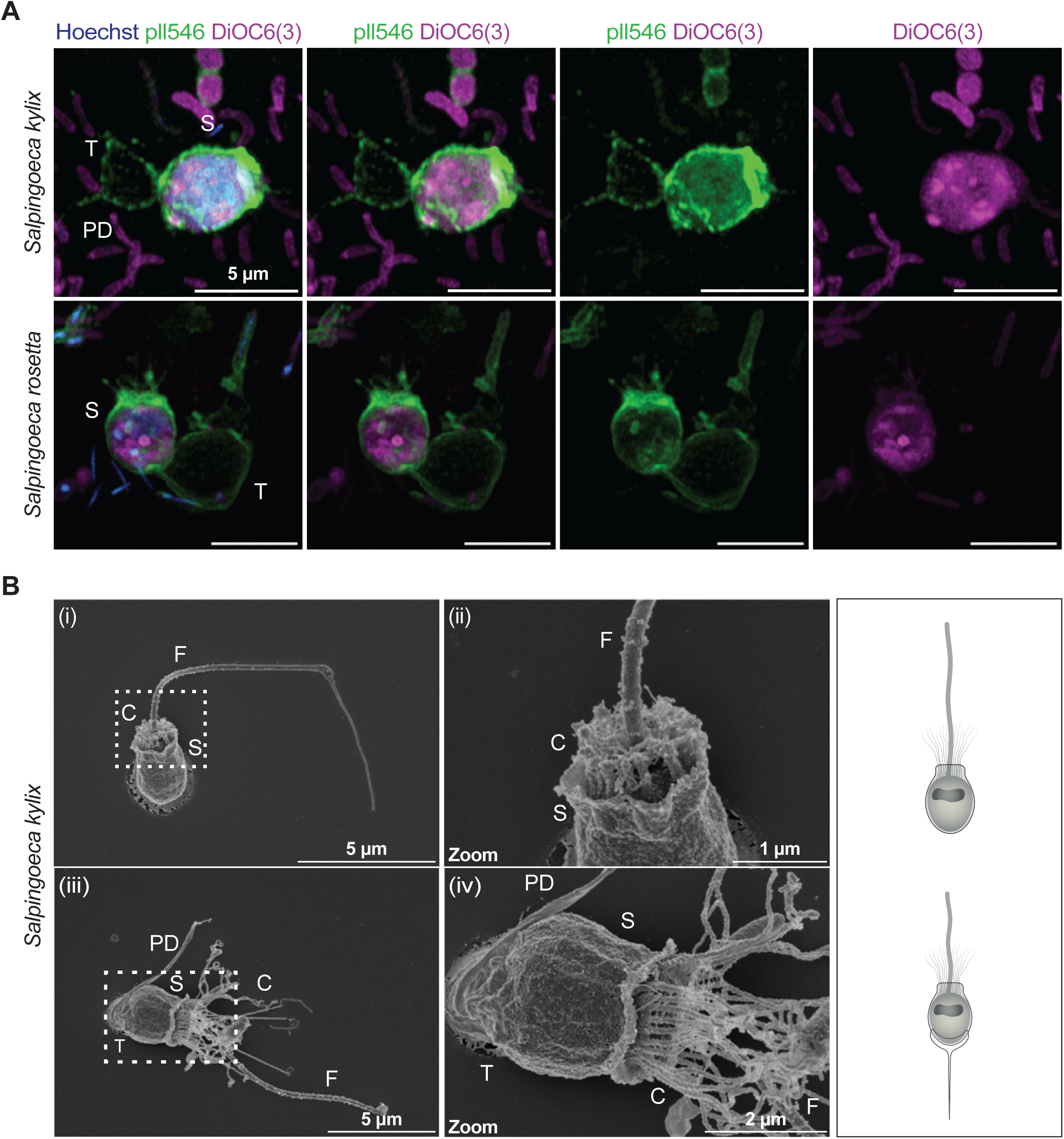
*Salpingoeca rosetta* and *Salpingoeca kylix* are characterized by the presence of an additional sheath. PD: pedicel, S: sheath, C: collar, F, flagellum, T: theca. **A)** Super-resolution microscopy of representative *Salpingoeca* cells abandoning their thecae, while retaining the additional outer sheath, stained with Hoechst (blue: DNA), PLL546 (green: extracellular structures) and DiOC6(3) (magenta: cell membranes). The newly isolated species are co-cultured with their bacterial prey, which are also stained. Scale bar: 5 μm. **B)** Scanning electron microscopy (SEM) images of *S. kylix* at distinct stages. (i) “Slow swimmer” cell showing an additional sheath, a short collar and a flagellum. White box indicates magnified regions. Scale bar: 5 μm. (ii) Magnified view of (i), where the sheath is seen partially covering the collar. Scale bar: 1 μm. (iii) Thecate cell displaying an additional sheath, theca, long collar, flagellum and pedice. White box indicates magnified region. Scale bar: 5 μm. (iv) Magnified view of (iii), highlighting the additional sheath. Scale bar: 2 μm.

**Supplementary Figure 3:**
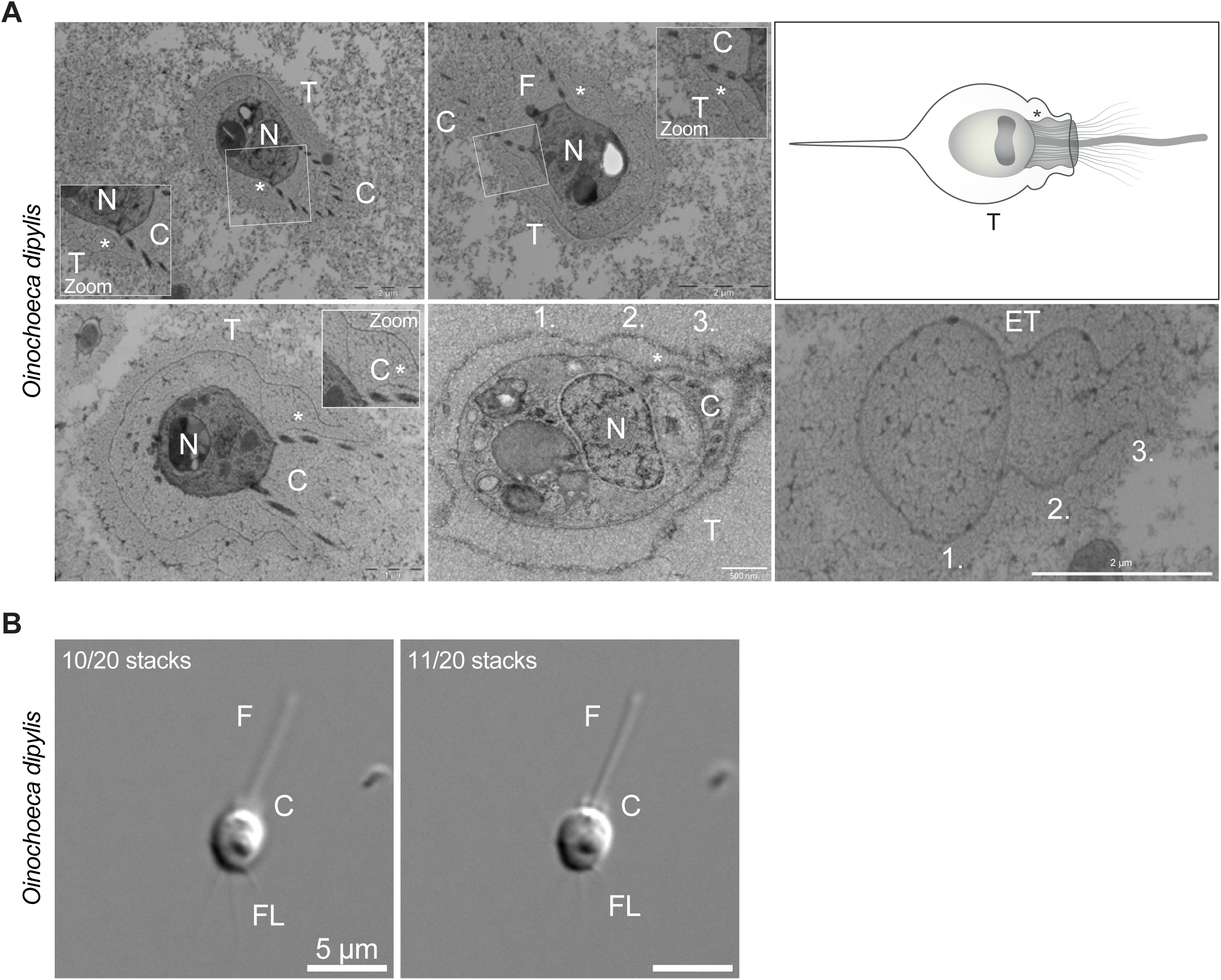
N: nucleus, F: flagellum, C: collar of microvilli, FL: filopodia and T: theca. **A)** Ultrastructural characterization of *Oinochoeca dipylis* via transmission electron microscopy (TEM). TEM sections of *O. dipylis* showing a thin theca that is separated from the cell body. Asterisks indicate the additional extracellular layer covering the collar of microvilli. The numbers 1, 2 and 3 indicate the three curved extensions of the theca. Scale bar: 2 μm, 1 μm and 500 nm. **B)** Differential interference contrast (DIC) microscopy images showing a crawling *O. dipylis* cell acquired at different focal planes to reveal different cellular features. The cell has a flagellum, a collar of short microvilli, filopodia and lacks a theca. Scale bar: 5 μm.

**Supplementary Figure 4:**
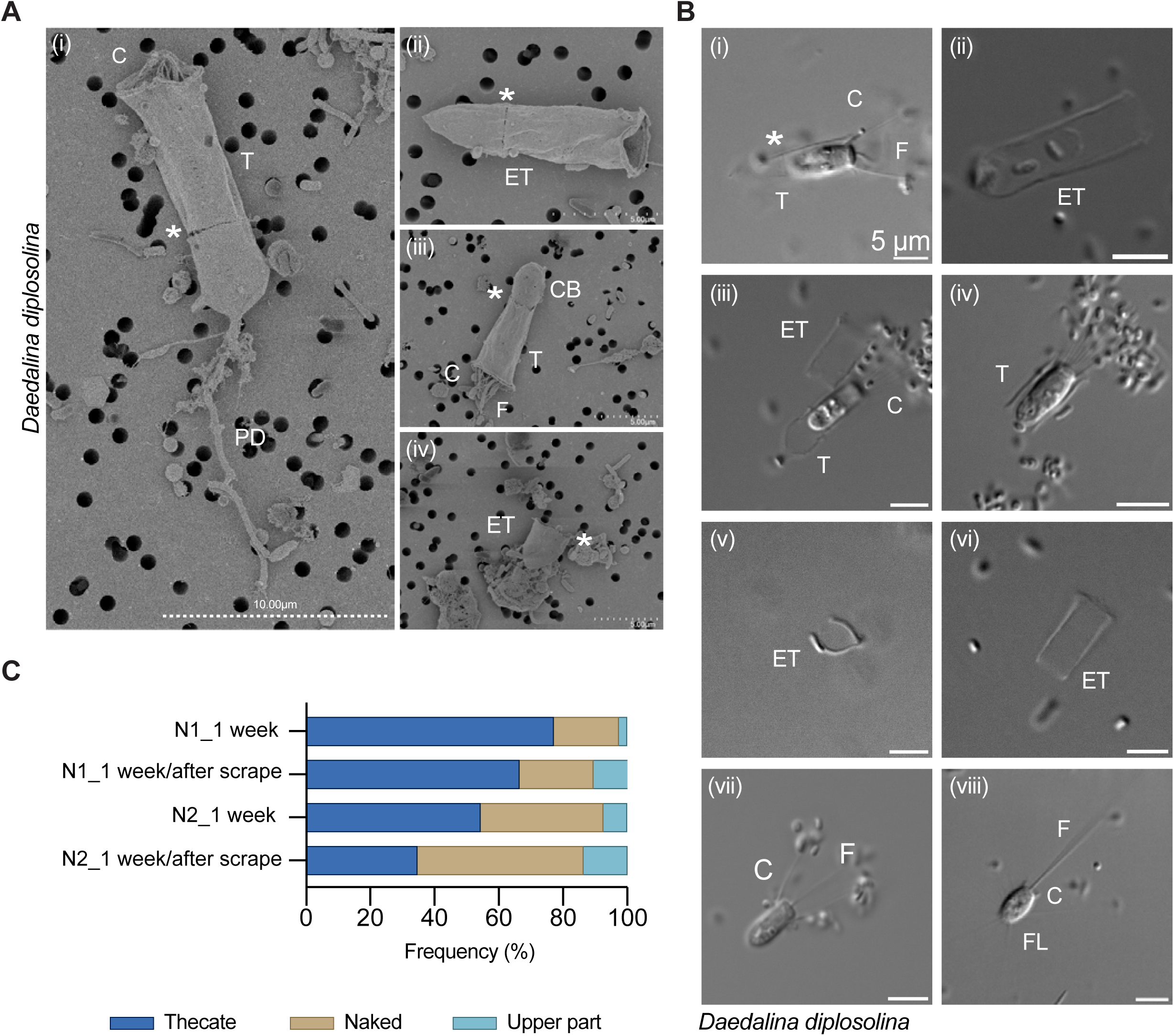
Morphological diversity and origin of the crawling cell phenotype in *Daedalina diplosolina*. T: theca, ET: empty theca, C: collar, F: flagellum, FL: filopodia. **A)** Scanning electron microscopy (SEM) reveals diverse morphotypes, including thecate cells with intact two-part thecae extending into a pedicel (i), empty thecae (ii), cells retaining only the upper thecal part (iii), and empty bottom thecal parts (iv). Asterisks mark the junction between the two parts of the theca. Scale bars: 10 μm (i) or 5 μm (ii-iv). **B)** DIC microscopy shows various life stages present following mechanical disruption, including intact thecate cells (i), empty thecae (ii, v-vi), cells with only one of the thecal parts (iii-iv), free-swimming naked cells (vii), and crawling cells with a short collar, long flagellum, and filopodia (viii). Scale bars: 5 μm. **C)** Morphotype proportions recorded one week after seeding and following mechanical disruption in two independent experiments, showing increased numbers of naked cells and cells with only the upper thecal part after mechanical stress. In the first experiment (N1), 153 cells were counted before scraping and 152 after scraping. In the second experiment (N2), 241 cells were counted before scraping and 242 after scraping.

**Supplementary Figure 5:**
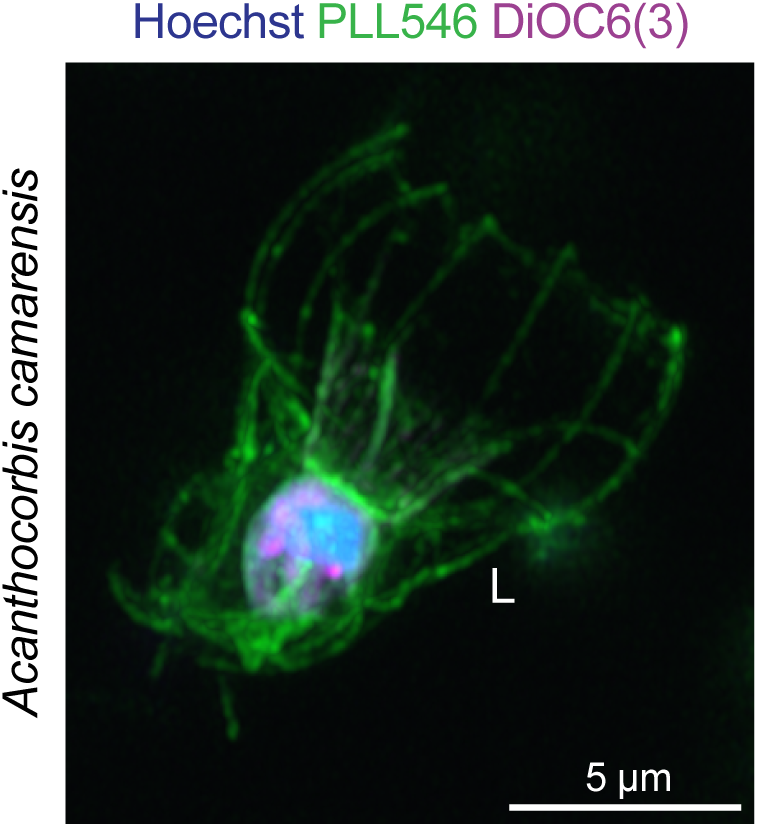
PLL546 clearly stains the lorica of *Acanthocorbis camarensis*. Super-resolution microscopy of *A. camarensis* stained with Hoechst (blue; DNA), PLL546 (green; extracellular structures), and DiOC6(3) (magenta; cell membranes), showing clear labelling of the lorica (L). Scale bar: 5 µm.

**Supplementary material** (available on FigShare at DOI: 10.6084/m9.figshare.34737594)

**Supplementary Table 1:** Taxa, data sources and GenBank accessions or assembly contigs used in the concatenated 18S, 5.8S and 28S rRNA gene phylogeny.

**Supplementary Material 1-14:** 3D reconstruction and 3D surface analysis videos of thecates or empty thecas stained with PLL546 (green) to highlight extracellular structures.

**Supplementary Material 15:** DIC video of a *Daedalina diplosolina* crawling cell settling and beginning to generate a theca.

**Supplementary Material 16:** DIC video of a thecate *Daedalina diplosolina* cell undergoing two back-to-back inverted divisions, each characterized by the emergence of a crawling cell. The second division is shown as the live-cell imaging sequence in Figure 7.

